# TGFβ-ANGPT2-Tie2 axis in cancer-associated fibroblasts reprograms oral cancer cells to embryonic-like cell state with predictive significance of poor prognosis

**DOI:** 10.1101/2024.06.29.601319

**Authors:** Paromita Mitra, Uday Saha, Kingsly Joshua Stephen, Priyanka Prasad, Ankit Kumar Patel, BV Harshvardhan, Santosh Kumar Mondal, Sillarine Kurkalang, Sumitava Roy, Arnab Ghosh, Shantanu Saha Roy, Jayasri Das Sarma, Nidhan Kumar Biswas, Moulinath Acharya, Rajeev Sharan, Pattatheyil Arun, Mohit Kumar Jolly, Arindam Maitra, Sandeep Singh

**Author notes:** Corresponding Address: Sandeep Singh, PhD Associate Professor National Institute of Biomedical Genomics Kalyani, WB 741251 India. AKP, currently affiliated with Umea University, Sweden. RS, currently affiliated with HCG Cancer Center, Kolkata, India. SK, currently affiliated with Comprehensive Cancer Center, University of Chicago medicine, IL, USA. **Conflict of interest statement:** The authors declare no potential conflicts of interest.

## Abstract

Myofibroblastic cancer-associated fibroblasts (CAFs) in tumor stroma is identified as poor-prognostic indicator in oral cancer; however, biological mechanisms are largely unexplored. Here, we discovered the role of autocrine or exogenous transforming growth factor beta (TGFβ) in inducing Tunica Interna Endothelial cell kinase 2 (Tie2) -signaling through histone deacetylase-mediated downregulation of Tie2-antagonist, Angiopoietin-2 in CAFs, responsible for induction and maintenance of myofibroblastic differentiation. To understand the influence of CAF-specific Tie2-signaling on cancer cell properties, we performed CAF-Cancer cell co-culture and its single-cell RNA sequencing (scRNA-Seq). Distinct clustering of CAFs suggested their transcriptional heterogeneity, driven by TGFβ-Tie2 activation. Interestingly, CAF-specific Tie2-signaling was responsible to reprogram cancer cells, producing embryonic-like cell state with increased stemness and EMT signatures. Importantly, both the Tie2-specific gene expression signature as well as reprogrammed cancer cell specific gene expression modules were validated respectively in fibroblasts clusters and malignant cell clusters in two independent earlier reported scRNAseq studies of HNSCC tumors. Highlighting the translatability of our study, the gene expression signature derived from reprogrammed cancer cells showed significant association with poor prognosis in HNSCC patient of TCGA cohort. Pharmacological inhibition of Tie2-signaling in CAFs, significantly abrogated the tumor initiating ability of co-cultured oral cancer cell lines. Overall, combining our molecular and computational analysis, we may propose Tie2 as a novel factor responsible for CAF mediated cancer cell plasticity, associated with aggressive nature of oral cancer.

**Teaser:** Tie2-signaling is activated in myofibroblasts which impacts the behaviour of malignant cells by inducing cancer cell plasticity to acquire stemness.

## Introduction

Cancer-associated fibroblasts (CAFs) in tumor microenvironment (TME) is known to undergo changes that often promote tumor growth and survival (1) In squamous cell carcinoma, stroma serves as a lifeline which provides essential nutrients, complex secretome of chemokines, cytokines and matrix forming (e.g. collagen) and degrading (e.g. matrix metalloproteinase) factors for tumor growth (2) and plays pivotal role in tumor metastasis (3). Thus, modulation of activity of CAFs to perturb their interactions with cancer cells have garnered attention and promised to innovate therapeutic strategies (4); however, targeted therapy against CAFs has remained to be elucidated. This may be primarily because CAFs are heterogeneous population which may play context dependent roles (5,6). Though CAFs are found to facilitate cancer progression, indiscriminate depletion of CAFs has also shown to promote tumor growth (7). Such paradoxical observation of CAFs-functions warrants deeper understanding about its biology (8,9).

Several lines of evidence have supported the notion that TGFβ produced in tumor microenvironment modulates adjacent fibroblasts into myofibroblasts (10–12) indicated by an increased expression of alpha smooth muscle actin (αSMA) and stress fibre formation (13,14). We have previously reported two diverse subtypes of CAFs; C1-CAFs (with lower score) and C2-CAFs (with higher score) of αSMA-stress fibre positive myofibroblast in oral cancer, where C2-CAFs supported higher stemness in cancer cells (15). Stemness is defined as the ability of cancer cells to display long-term regeneration ability, giving rise to heterogeneous subpopulations of cancer cells and linked with several critical aspects including cancer initiation, progression and treatment responses (16–18). Targeting these stem-like cancer cells (SLCCs) may be crucial to prevent relapse and for overall success of treatments (19,20).

Tunica interna endothelial cell kinase 2 (Tie2) gene, also known as TEK or angiopoietin-1 receptor, encodes for a receptor tyrosine kinase. Substantial reports have supported the potential antagonistic role of ANGPT2 on Tie2 signalling, while ANGPT1 acts as agonist (21,22). Studies on Angiopoietin/Tie2 pathway have been majorly focused on endothelial cell (EC) functions, related to vessel maturation and vascular integrity (23,24). Increasing literature have gathered evidences of Tie2-activation in pericytes, macrophages and hematopoietic stem cells, as well (25–27). Role of Tie2 in cancer tissue is reported in breast tumor-bone microenvironment, where Tie2-positive myeloid cells were found to be involved in osteoclast differentiation and osteolytic bone invasion of murine breast cancer cell line (28). Also, elevated ANGPT1/Tie2 signaling was positively correlated with increased cell proliferation and migration in thyroid carcinoma (29). Tie2-positive cervical cancer cells are recently reported to induce VEGFR2 and Tie2 expression in endothelial cells and promote angiogenesis (30). Moreover, Tie2-expressing cervical cancer cell-derived exosomes transports Tie2 protein in infiltered macrophages, and increase angiogenesis (31). Similarly, neovascular endothelial cells showed higher expression of Tie2 in hepatocellular carcinoma (32). Tie2 expression in oral tumor tissues is studied briefly (33). Moreover, Tie2 was also among the top upregulated genes in patient derived C2-CAFs in our earlier report (15); however, its fibroblasts specific expression and precise role in the biology of oral tumor progression has remained to be elucidated.

Here, we report that myofibroblastic differentiation by TGFβ facilitated C1-CAFs to acquire transcriptional changes and functions similar to C2-CAFs. Importantly, we found upregulation of Tie2 expression in TGFβ-induced as well as patient derived constitutive C2-CAFs, where Tie2 activity was found essential in initiation and maintenance of TGFβ-induced myofibroblastic differentiation and acquiring properties of C2-CAFs. Furthermore, Tie2-signal in activated-CAFs was responsible for reprogramming oral cancer cells to acquire embryonal gene expression state with increased stemness and epithelial to mesenchymal transition (EMT). Moreover, validating our *in vitro* results, similar CAF-induced cancer cell reprogramming was also identified in HNSCC tumors at single cell level, and found associated with poor prognosis in TCGA-HNSCC patient cohort, suggesting the clinical implication of our study. Targeting Tie2-activity in oral-CAFs led to reduced tumorigenic ability of cancer cells; demonstrating wider applicability of Tie2, beyond endothelial cell specific functions.

## RESULTS

### C2-CAFs expressed higher levels of Tie2 and positively correlated with αSMA-high stromal fibroblasts in primary tumors

Provided that TGFβ induces myofibroblastic differentiation, and based on our previous study where C2-CAFs demonstrated myofibroblastic phenotype; we first tested if C1-CAFs may acquire status of C2-CAFs upon TGFβ induction. Interestingly, stimulation of TGFβ (10 ng/ml) led to a significant increase in frequency of cells having αSMA-positive stress fibers in all three tested, patient-derived C1-CAFs; indicative of myofibroblastic differentiation (Figure S1A). Moreover, TGFβ-induction resulted in gain of C2-CAFs associated genes (*FN1, SERPINE1, ITGB1*); while, genes associated with C1-CAFs state (*FOXF1, EYA1, RUNX2*) were downregulated compared to untreated control, suggesting TGFβ-induced transition of C1-CAF to C2-CAF status (Figure S1B). αSMA is associated with contractile apparatus of smooth muscle cells and myofibroblasts and exhibits matrix remodelling ability (34). Notably, TGFβ- induced CAFs had better matrix remodelling ability than untreated C1-CAF group (Figure S1C, i-ii). Taken together, TGFβ-induction clearly converted C1-CAFs (αSMA^low^) to C2-CAFs (αSMA^high^). For ease of understanding we have labelled C1-CAFs as UT-CAF and TGFβ-induced C1-CAFs as TGF-CAF.

To explore tumor-stroma interaction, UT-CAF or TGF-CAF were co-cultured with cancer cells. Following co-culture, cells were separated using FACS and bulk-RNAseq was performed on sorted cells, subsequently (Figure 1A). We found that 886 and 1065 genes were upregulated and downregulated respectively (log2FC>1, pvalue ≤0.05) in TGF-CAF compared with UT-CAF (Table S1). Gene set enrichment (35), with Cytoscape analyses suggested enrichment of key regulatory pathways involving RTKs, PI3K/AKT, focal adhesion, JAK-STAT pathway, cytokines- and interleukins-mediated pathways (Table S2) in TGF-CAF (Figure S2A- D). Receptor tyrosine kinases (RTKs) are key regulatory trans-membrane receptors which made them suitable candidates for therapeutic target (36). Activation of RTK leads to downstream activation of MAPK and PI3K-AKT pathway. With the aim to identify common regulators of these pathways; *TEK* (*Tie2)*, *ERBB3*, *FGFR2*, *EREG*, *TGFA*, *FGF5*, *MET*, and *FGF2* were commonly upregulated in TGF-CAF (Figure 1B, S2E), with Tie2 being the most upregulated RTK in TGF-CAF. Also, genes associated with Tie2 signaling were significantly enriched in TGF-CAF (Figure S2F-i) and Tie2 upregulation was verified by qPCR (Figure S2F-ii). This collectively prompted us to explore the expression and function of Tie2 in oral-CAFs in response to TGFβ.

**Figure 1:**
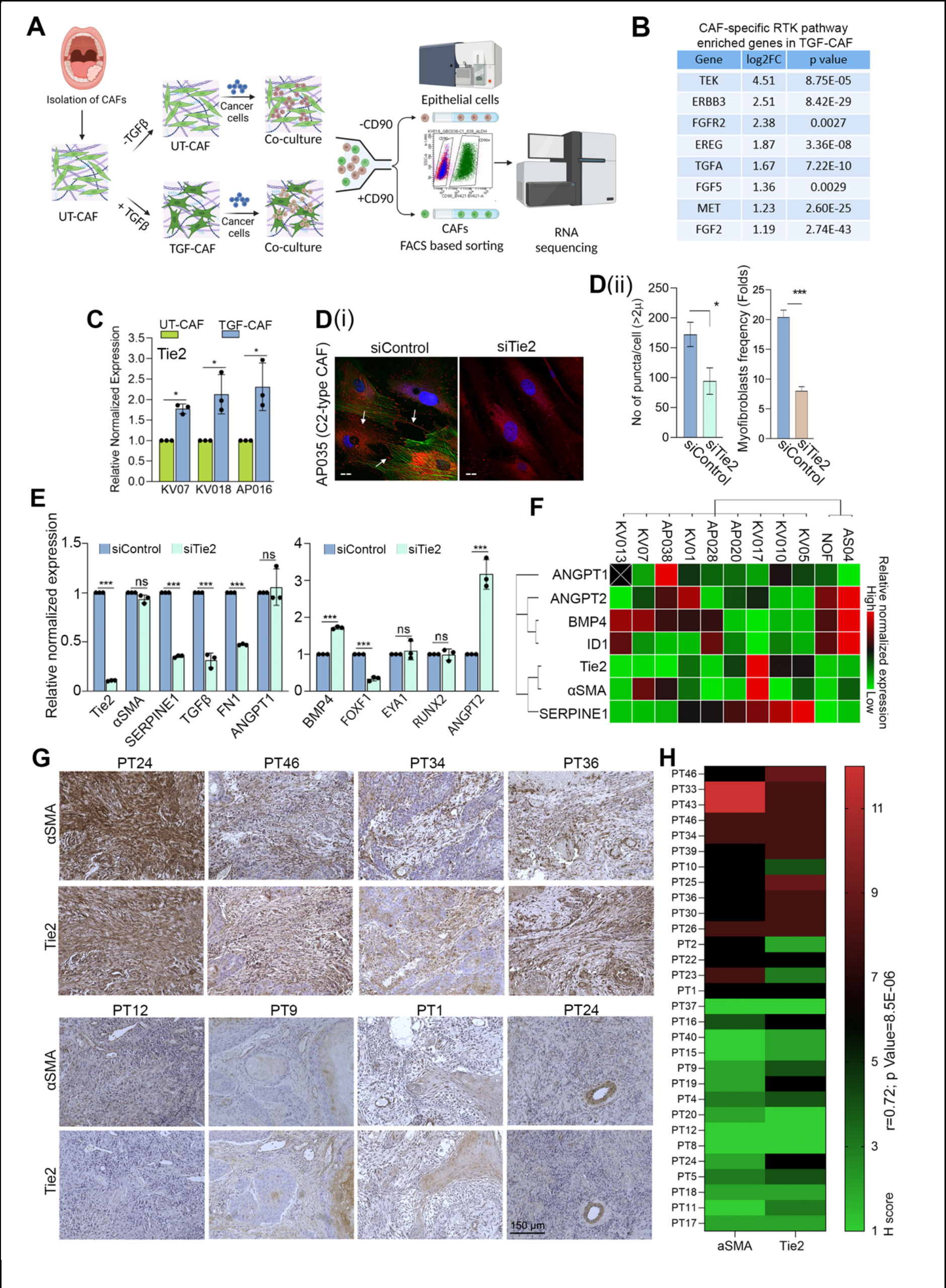
C2-CAFs expressed higher levels of Tie2 and positively correlated with αSMA- high stromal fibroblasts in primary tumors: (A) Schematic depicting experimental design for co-culture of UT-CAF and TGF-CAF along with cancer cells and downstream processing. **(B)** List of eight common upregulated genes between RTK, PI3K, MAPK in TGF-CAF. **(C)** qPCR analysis of *Tie2* in three different primary CAFs under untreated (UT-CAF) or 10 ng/ml TGFβ- induced (TFG-CAF) conditions. **(D)** (i) Images of constitutively activated C2-CAF (AP035), stained for *a*SMA (green), pTie2 (Y992) (Red), nucleus (DAPI, purple) after RNAi mediated silencing of Tie2 (siTie2). Scrambled siRNA (siControl) was used as a control. Arrowhead indicates pTie2 (Y992) positive puncta. (ii) frequency of CAFs with myofibroblast-phenotype (with *a*SMA- positive stress fibre) and pTie2 (Y992) puncta was quantified using ImageJ. Scale bars, 20 µm. **(E)** qPCR analysis of C1-CAF related genes (*BMP4, EYA1, RUNX2, FOXF1, ANGPT2*) and C2-CAF related genes (*Tie2, TGFꞵ, SERPINE1, aSMA, FN1, ANGPT1*) in constitutively activated C2-CAF following Tie2 knock-down. **(F)** Heatmap showing qPCR-based expression of C1- and C2- CAF related genes across different primary CAFs from oral cancer patients and normal oral fibroblasts. **(G)** Representative images of human oral tumor tissues detected for αSMA and Tie2 protein expression using IHC. **(H)** Heatmap showing correlation between H-score of αSMA and Tie2 protein in oral tumor stroma. Heatmap showing correlation between H-score of αSMA and Tie2 protein in tumor stroma. Scale bars, 150 µm. *P<0.05, **P<0.01, ***P<0.001.

First, a direct upregulation of Tie2 in TGFβ-induced condition, independent of co- culture with cancer cell was observed in all tested CAFs (Figure 1C). Thus, we explored the Tie2 association with myofibroblastic phenotype and maintenance of C2-like state of CAFs. Interestingly, silencing of Tie2 in patient derived C2-CAFs showed significantly reduced myofibroblasts frequency, compared to control (Figure 1D, i-ii). Reduced Tie2- phosphorylation confirmed downregulation of Tie2-activity upon Tie2-silencing, depicted by loss of puncta of phosphorylated Tie2 (Y992) and mature focal adhesions with length 2μm or longer (37). Importantly, Tie2 silencing in C2-CAFs also showed concomitant downregulation of tested C2-CAF related genes (*SERPINE1, FN1, TGF*β) whereas, C1-CAF related gene *BMP4* was upregulated. Additionally, antagonist ANGPT2 was upregulated in Tie2 silenced C2-CAFs without having any effect on its agonist, ANGPT1 (Figure 1E). Therefore, to substantiate, we further explored this correlation in ten different oral tumor derived CAFs and one normal oral mucosal fibroblast (NOF) (Figure 1F). While ANGPT1 expression did not specifically associate with any specific gene; interestingly, gene expression based unsupervised clustering grouped *Tie2* with *αSMA* and *SERPINE1*, whereas its antagonist *ANGPT2* clustered with *BMP4* and its downstream gene *ID1*. Collectively, results established a strong correlation between Tie2 expression with myofibroblastic C2-like state of CAFs. Encouraged from the results, we evaluated the tumor stromal expression of αSMA and Tie2 on serial sections of surgically resected human oral tumor tissues (n=30) (Figure 1G). To our interest, we observed significantly higher H-score of Tie2 in tumors having αSMA-high stromal fibroblasts as compared to tissues having αSMA-low stroma (Figure 1H). Thus, we obtained strong support for Tie2 expression in myofibroblastic CAFs in tumor stroma.

### Tie2 plays an essential role in induction as well as sustenance of TGFβ-induced myofibroblastic differentiation of CAFs

With a hypothesis that Tie2 activity is required for TGFβ-induced myofibroblastic differentiation, we used a commercially available small molecule inhibitor, selective against Tie2 kinase (Tie2i) (38). Similar to our observation with Tie2 silencing, one hour pre-treatment with Tie2i before TGFβ induction showed significantly less frequency of myofibroblasts compared to DMSO control (Figure 2A) in two different patient-derived C1-CAFs. More importantly; even after CAFs were successfully induced to myofibroblasts by TGFβ, Tie2i effectively reversed this myofibroblast phenotype (Figure 2A) and downregulated C2-CAF associated genes *αSMA, SERPINE1* and *Tie2* (Figure S3A, S3B).

**Figure 2:**
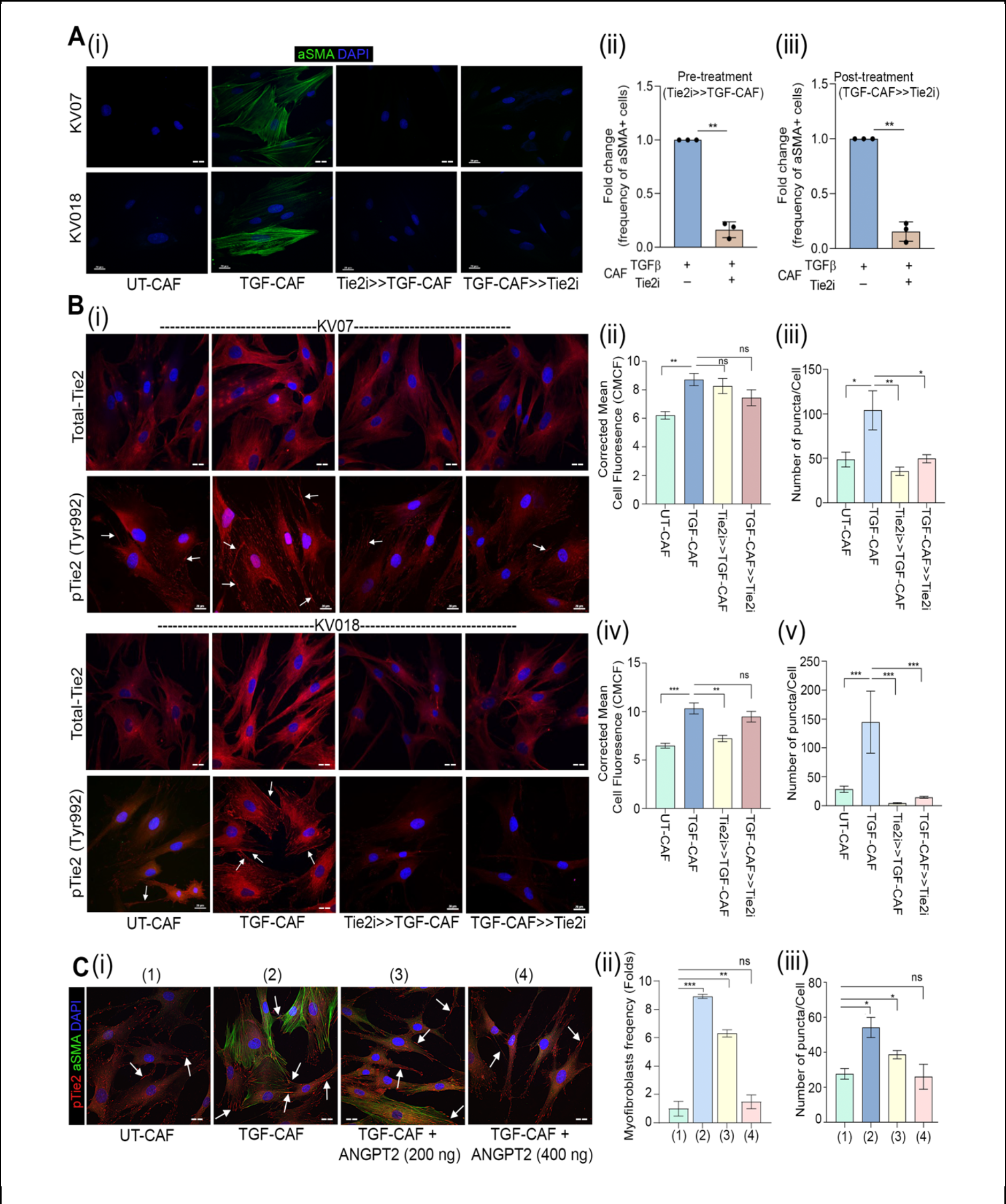
Tie2 plays essential role in induction as well as sustenance of TGFβ-induced myofibroblastic differentiation of CAFs. (**A)** (i) Representative images and quantification of myofibroblasts frequency in UT-CAF and TGF-CAF. Tie2-inhibitor was added 1 hr before TGFβ induction (Tie2i>>TGF-CAF) or 48 hrs after TGFβ induction (TGF-CAF>>Tie2i). Cells were quantified using ImageJ. (ii-iii) Frequency of αSMA stress-positive cells were plotted for three different patient derived CAFs. **(B)** (i) Representative images of Tie2 and pTie2 (Y992) in UT-CAF and TGF-CAF. or afterTie2 inhibition was done 1 hr before (Tie2i>>TGF- CAF) or for 6 hrs after TGFβ induction (TGF-CAF>>Tie2i). (ii-v) Bar graph showing quantification of total Tie2 protein and pTie2 (Y992) puncta, calculated using ImageJ software. **(C)** (i) Representative images of αSMA and pTie2 (Y992) in UT-CAF, TGF-CAF or with increasing doses of ANGPT2 (200 ng/ml, 400 ng/ml) in presence of TGFb. Arrowhead indicates pTie2 (Y992) puncta. (ii) Bar graph showing cell frequency with αSMA stress-positive CAFs and (iii) pTie2 (Y992) expression by CAFs in given conditions. *P<0.05, **P<0.01, ***P<0.001. Scale bars, 20 μm.

Further, upon TGFβ-induction a significant increase in total-Tie2 protein and frequency of phosphorylated-Tie2 (Y992) puncta (Figure S3C (i, ii)) were observed for tested CAFs (Figure 2B, i-v). Importantly, one hour pre-treatment with Tie2i before TGFβ induction as well as six hour of Tie2-inhibition after complete myofibroblastic differentiation by TGFβ (post-treatment), both conditions showed reduced number of Tie2-phosphorylated puncta. Since, ANGPT2 is a known antagonist of Tie2-receptor activation, we next used soluble ANGPT2 to inhibit Tie2 signaling. Very interestingly, similar to Tie2i, reduced frequency of myofibroblasts (Figure 2C, i-ii) and number of Tie2-phoshorylated puncta (Figure 2C-iii) was observed after ANGPT2 addition. Taken together, results provided novel insights, where CAF-specific Tie2 activity was responsible for induction and maintenance of TGFβ-induced myofibroblastic phenotype as well as transcriptional state of C2-CAFs.

### Tie2-activity is regulated in an autocrine manner

To explore the mechanism behind TGFβ-induced Tie2-activation, we next used pharmacological inhibitor of these regulators directly on a patient-derived C2-CAFs (AP035), having constitutive-myofibroblastic phenotype (Figure 3A). As anticipated, Galunisertib (TGFβi) or Tie2i independently led to reduction in frequency of Phospho-Tie2 (Y992) positive puncta as well as myofibroblast frequency, as compared to control (Figure 3B, i,ii); suggesting endogenous TGFβ-mediated constitutive activation of Tie2 in C2-CAFs. More interestingly, both Tie2- and TGFβ-inhibited C2-CAFs showed significant downregulation of genes associated with C2-CAFs (*αSMA* and *SERPINE1*) with concomitant upregulation of genes associated with C1-CAFs (*BMP4* and *ANGPT2)* (Figure 3C, i-iii); indicating a transition of C2- CAFs, back to C1-CAFs.

**Figure 3:**
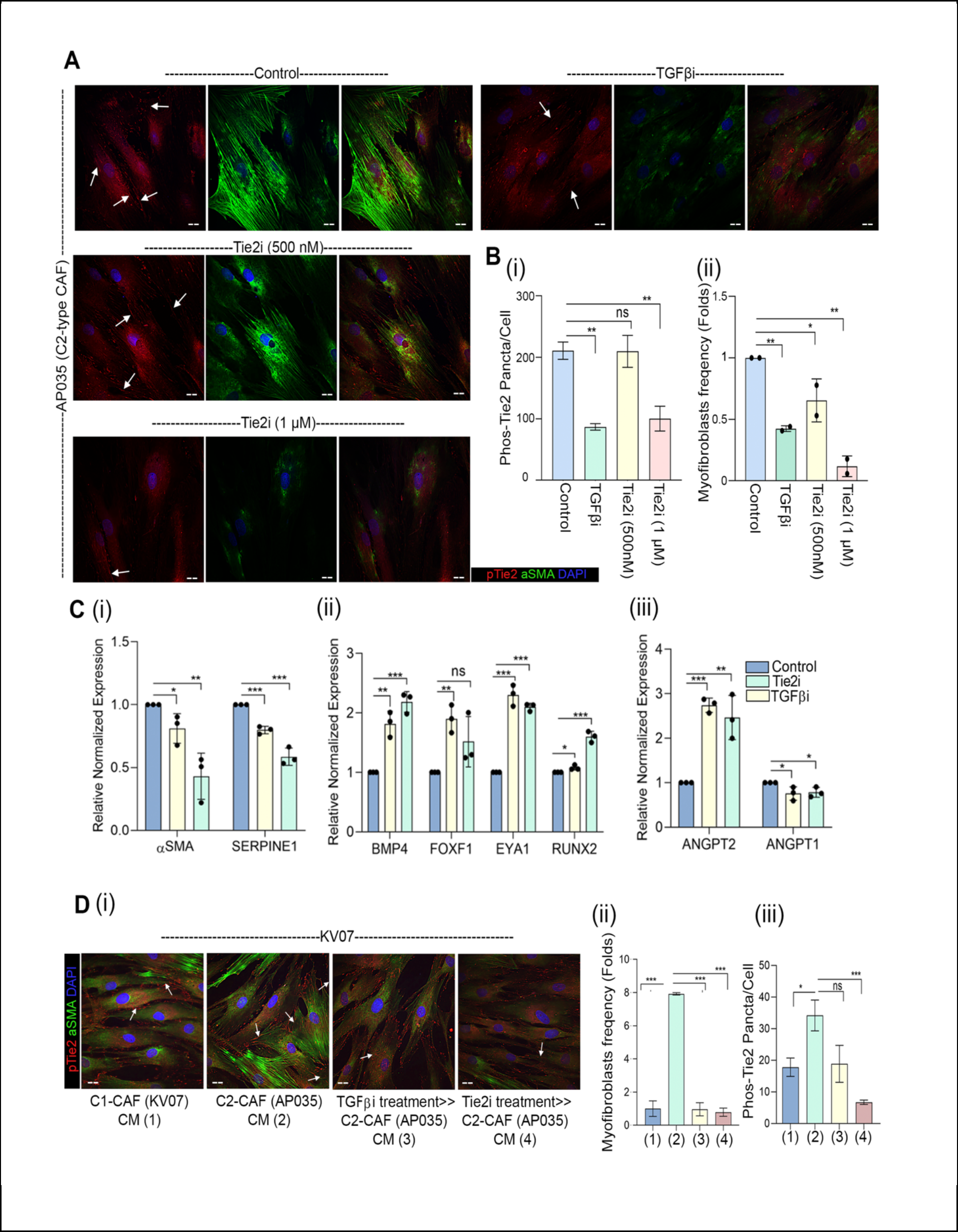
Tie2-activity is regulated in an autocrine manner. (A) Representative images of constitutively active C2-CAF (AP035) detected for αSMA and pTie2 (Y992) protein. Increasing doses of Tie2 inhibitor and TGFβ inhibitor (LY2157299; 1µM) were used to block respective receptor activity. Cells were quantified using ImageJ software. **(B)** (i) quantification of pTie2 (Y992) puncta and (ii) myofibroblasts frequency under these conditions. **(C)** qPCR analysis of (i) C2-CAF related genes (*SERPINE1*, *aSMA*), (ii) C1-CAF related genes (*BMP4*, *EYA1*, *RUNX2*, *FOXF1*), and (iii) ligand of Tie2 receptor (*ANGPT1*, *ANGPT2*) following Tie2 inhibitor and TGFβ inhibitor treatment in constitutively activated C2-CAF. Unstimulated CAF in same media was used as control. **(D)** (i) Representative images of C1-CAF (KV07) exposed to conditioned media from C1-CAF (KV07), C2-CAF (AP035), TGFβ inhibited C2-CAF (TGFβi>C2 CAF), Tie2 inhibited C2-CAF Tie2i>C2 CAF, for 48 hrs, detected for αSMA and pTie2 (Y992) (ii) myofibroblasts frequency and (iii) pTie2 (Y992) puncta was quantified using ImageJ. Arrowhead indicates pTie2 (Y992) puncta. *P<0.05, **P<0.01, ***P<0.001. Scale bars, 20μm.

Since both TGFβ and Tie2 signaling are activated through receptor-ligand interactions, we explored if secreted factors from C2-CAFs may act as drivers for acquiring and maintaining C2-CAF-like state. Conditioned media of KV07 (C1-CAFs) and AP035 (C2-CAFs) were collected and put over KV07 (C1-CAFs) (Figure 3D-i). Interestingly, conditioned media of C2-CAFs was sufficient to increase both, myofibroblasts frequency (Figure 3D-ii) and number of Phos-Tie2 (Y992)-positive puncta in C1-CAFs (Figure 3D-iii). This was significantly reduced when C1-CAFs were exposed to conditioned media collected from TGFβ-inhibited or Tie2-inhibited C2-CAFs (Figure 3D-ii, iii), suggesting the transition of C1-to C2-CAFs and maintenance of C2-CAF-like state in an autocrine manner, through secretory factors.

### TGFβ-induced histone deacetylation drives transcriptional state change associated with C1- to C2-CAF transition

To delve into the mechanisms further, we performed TGFβ induced gene expression analysis in a timeseries manner (Figure 4A). Suggesting the activation of TGFβ-signal, increased expression of *SERPINE1* was observed at as early as 6 hours which was maximum at 12 hours. Expression of endogenous *TGFβ* and *Tie2* genes also showed its peak levels by 12 hours of TGFβ-induction. *αSMA* gene showed upregulation only after 48 hours, interestingly; suggesting that Tie2 expression preceded *αSMA*-upregulation during myofibroblastic differentiation by *TGFβ*. While genes associated with C2-CAFs showed upregulation; we observed very sharp downregulation of C1-CAF specific genes *BMP4* and *ANGPT2* from as early as 6 hours. While, antagonist *ANGPT2* showed sustained downregulation for entire test-period (96 hours) of TGFβ induction, agonist *ANGPT1* was upregulated at later time point. Overall, these results indicated the presence of TGFβ-induced feed-forward loop of Tie2- activation by rapid suppression of ANGPT2 followed by upregulation of endogenous *TGFβ*, *Tie2* and *ANGPT1*. Since, addition of ANGPT2 was sufficient to block TGFβ-induced myofibroblastic differentiation (Figure 2C); thus, rapid suppression of ANGPT2 may be one of the most crucial events in TGFβ-induced transition of C1-CAFs into C2-CAF. Thus, we next performed chromatin immunoprecipitation to evaluate activation-marks using H3K27- acetylation for ANGPT2 and BMP4 locus. Interestingly we observed reduced H3K27- acetylation on TATA binding site (-1600 bp) and initiator site (-400 bp) of ANGPT2 promoter and the tested locus of *BMP4* promoter (-708 bp) in TGF-CAF, compared to UT-CAF (Figure 4B). Next, using three different C1-CAFs, we tested the effect of TGFβ-induction in presence of potent histone deacetylase (HDAC) inhibitor, Valproic acid (VPA). Suppressive effect of TGFβ on all tested C1-CAFs associated genes, *BMP4, EYA1, FOXF1, RUNX2* and *ANGPT2* were significantly much lower, in presence VPA (Figure 4C). Thus, data strongly suggested the pivotal role of TGFβ-induced deacetylation of open chromatin on C1-CAF associated genes during myofibroblastic differentiation leading to the transition to C2-CAFs (Figure 4D).

**Figure 4:**
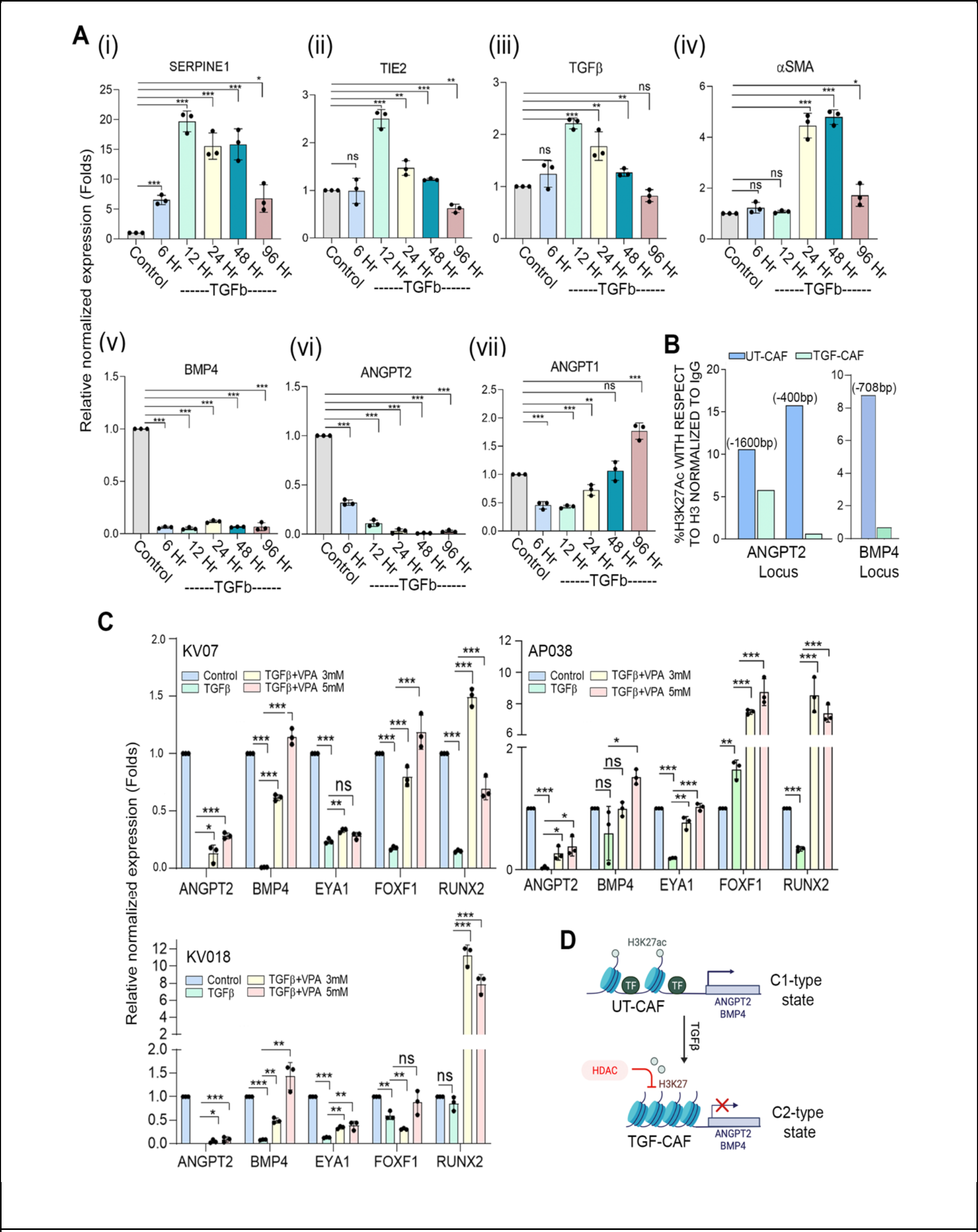
TGFβ-induced histone deacetylation drives transcriptional state changes associated with transition of C1- to C2-CAFs: (A) (i-vii) qPCR analysis of *SERPINE1, aSMA, TGFꞵ, Tie2, BMP4, ANGPT2* and *ANGPT1* in C1-CAF following 10 ng/ml TGFβ stimulation in time dependent manner as indicated. Relative abundance of mRNA was normalized with unstimulated CAF (Control) of respective time points. **(B)** Chromatin immunoprecipitation analysis of H3K27-acetylation status on ANGPT2 (TATA binding site -1600 bp; initiator site -400 bp) and BMP4 promoter (-708 bp) in C1-CAF with or without 10 ng/ml TGFβ. Unstimulated CAFs were used as control. **(C)** qPCR analysis showing expression of C1-CAF related genes, *BMP4*, *EYA1*, *RUNX2*, *FOXF1* and *ANGPT2* with or without valproic acid (3mM, 5mM) in presence of 10 ng/ml TGFβ. Unstimulated CAFs were used as control. D. Schematic model of HDAC-mediated suppression of C1-CAF related genes. *P<0.05, **P<0.01, ***P<0.001.

### Endogenous TGFβ is necessary and sufficient for driving Tie2-ANGPT signalling

Since, TGFβ-induced suppression of ANGPT2 was found important for Tie2- phosphorylation and myofibroblastic differentiation, we next explored if downregulating endogenous ANGPT2 may be sufficient for Tie2-phosphorylation in oral CAFs. Interestingly, ANGPT2 silencing in C1-CAF, did not result in any significant change in pTie2 (Y992) positive puncta (Figure 5A); however, addition of ANGPT1 increased the number of phosphorylated- Tie2 puncta in ANGPT2-silenced C1-CAFs (Figure 5B-C). Thus, downregulation of ANGPT2 was necessary but may not be sufficient for Tie2 phosphorylation in oral CAFs. Since, TGFβ- induced CAFs showed upregulation of endogenous-TGFβ, Tie2 and ANGPT1 along with suppression of ANGPT2, we next silenced increased levels of TGFβ, Tie2 or ANGPT1 in TGF- CAF. As anticipated, Tie2 and ANGPT1 silencing resulted in lowering in pTie2 (Y992) and myofibroblasts frequency in TGF-CAF (Figure 5D, 5E). Interestingly, even silencing of upregulated endogenous-TGFβ also suppressed Tie2-phosphotylation; supporting the role of endogenous-TGFβ in maintaining myofibroblast phenotype. Silencing of respective genes was confirmed by RT-PCR (Figure 5F). Importantly, reduced levels of all three genes showed increased expression of ANGPT2 in TGF-CAF, suggesting it to be in an interconnected regulatory signaling loop (Figure 5F).

**Figure 5:**
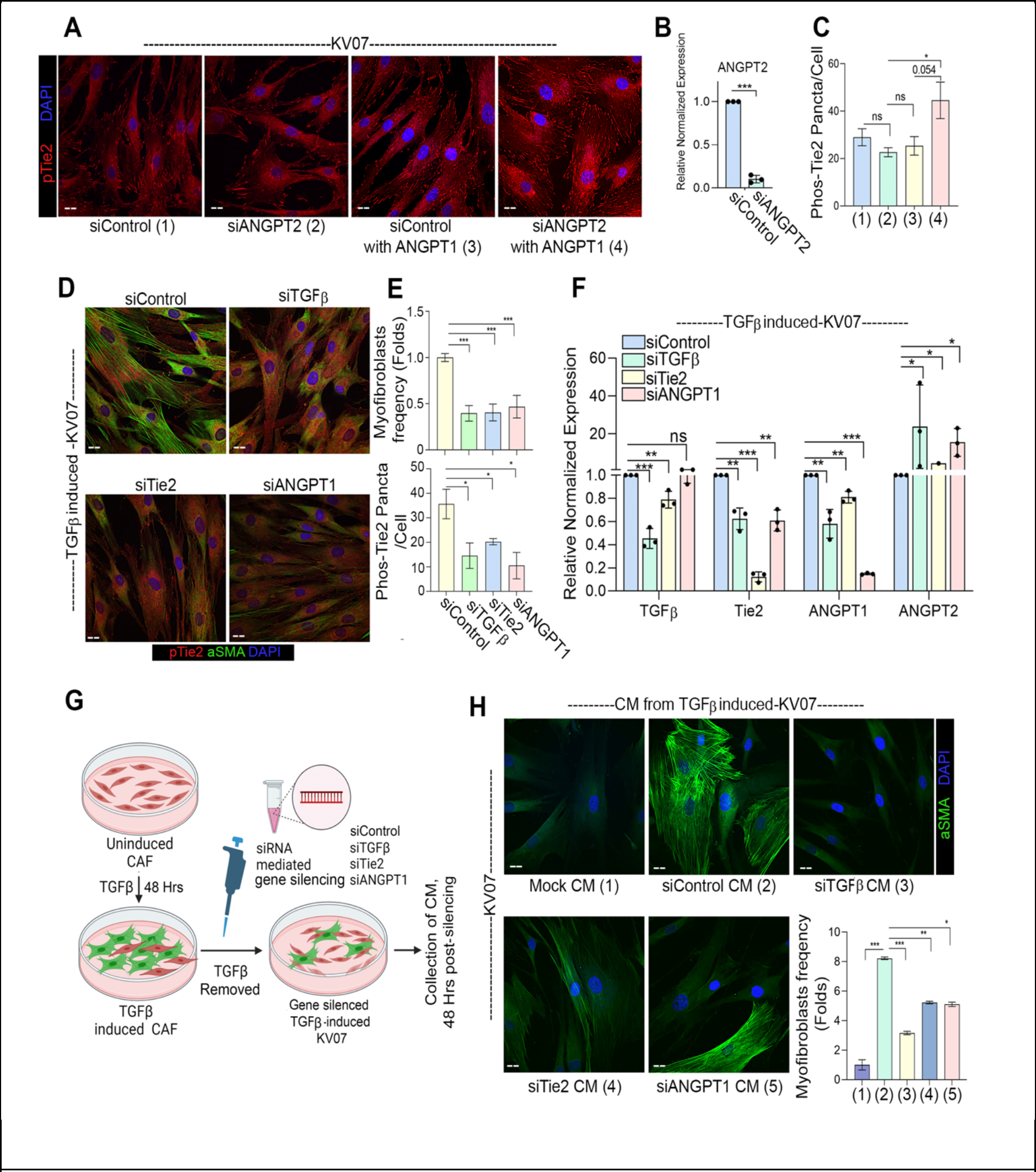
Endogenous-TGFβ is necessary and sufficient in driving Tie2-ANGPT signaling. **(A)** Representative images of ANGPT2 silenced C1-CAF with or without ANGPT1 stimulation (400 ng/ml) for 6 hrs, detected for pTie2 (Y992) protein. Scrambled siRNA was used as control. **(B)** qPCR analysis of *ANGPT2* following *ANGPT2* knockdown in C1 CAF. **(C)** Quantification of pTie2 (Y992) puncta using ImageJ. **(D)** Representative images of αSMA and pTie2 (Y992) protein detected by immunofluorescence staining, upon gene silencing of TGFβ, Tie2 and ANGPT1 in TGF-CAF. Scrambled siRNA was used as a control. **(E)** Myofibroblasts frequency and pTie2 (Y992) puncta was quantified using ImageJ. **(F)** qPCR analysis of *TGF*β*, Tie2, ANGPT1* and *ANGPT2* followed by knockdown of *TGF*β*, Tie2* and *ANGPT1* in TGF-CAF. **(G)** Schematic model suggesting experimental design of conditioned media (CM) collection from TGF-CAF following *TGF*β, *Tie2* and *ANGPT1* gene knockdown. **(H)** Representative images showing myofibroblasts frequency in uninduced C1-CAF exposed to the CM collected from TGF-CAF after TGFβ, Tie2 or ANGPT1 gene-silencing. C1- CAF exposed to C1-CAF CM was used as control. Myofibroblasts frequency was quantified using ImageJ. *P<0.05, **P<0.01, ***P<0.001. Scale bars, 20μm.

Since, there was an increase in ANGPT2 expression after silencing of induced levels of *TGFβ, Tie2* or *ANGPT1* in TGF-CAF, this prompted us to test if conditioned media (CM) from these experiments may have reduced myofibroblastic differentiation potency (Figure 5G). As anticipated, conditioned media from control-siRNA transfected TGF-CAF was sufficient to significantly increase the myofibroblasts frequency in UT-CAF. However, CM collected from *TGFβ, Tie2* or *ANGPT1* siRNA transfected TGF-CAF showed significantly lower frequency of myofibroblasts (Figure 5H). Collectively, these results led us to conclude that either extrinsic or endogenous-TGFβ in oral CAFs led to the activation of Tie2-ANGPT signal; responsible for transition of C1-CAFs to C2-CAFs and acquiring myofibroblast phenotype.

### C1-CAFs or C2-CAFs derived gene expression signatures showed concordance with patient derived BMP4-High and ITGA3-High CAFs respectively

To get deeper understanding about the intricate interplay between different CAF- phenotypes and their influence on oral cancer cells, we performed single cell RNA sequencing for co-cultures of oral cancer cells with C1-CAFs (UT-CAF) or TGFβ-induced CAFs (TGF-CAF) or TGFβ-induced-Tie2-inhibited (TGF>>Tie2i-CAF), separately (Figure S4A, B (i,ii,iii)). Based on module scores of canonical markers of epithelial- and CAFs- gene-sets (39), we identified CAF clusters from each of the co-culture conditions, having high-scores for CAF gene-set and low score for cytokeratin enriched epithelial gene set (Figure S5A, 6A). Unsupervised re-clustering of segregated 11391 CAFs from 3 different conditions (Figure 6B) showed transcriptional divergence on UMAP projection between UT-CAF and TGF-CAF and TGF>>Tie2i-CAF acquiring distinct transcriptional states. ‘Pseudo-time analysis’ performed using ‘Monocle3’, demonstrated the origin of TGF-CAF from UT-CAF at significant scale and depth; whereas, TGF>>Tie2i-CAF displayed a retrogressive transcriptional behaviour to remain in middlemost part of trajectory indicating a reversal of TGF-CAF towards UT-CAF upon Tie2-inhibition (Figure 6C (i, ii), S5B). Clusters belonging to UT-CAF such as 10,8,11,13 had a lower pseudo- time value than that of clusters comprises of TGF-CAF and TGF>>Tie2i-CAF, depicting a continuous evolution of CAF phenotypes from UT CAF to TGF-CAF through TGF>>Tie2i-CAF (Figure S5B). Next, we performed pseudo-bulk analysis of scRNAseq data to evaluate the cell-state specific combined features; where individual UT-CAF (Red), TGF-CAF (Green) and TGF>>Tie2i-CAF (Blue) were computed for ‘AUCell scores’ for TGFβ- or Tie2-signaling associated gene-sets as signatures. Interestingly, it showed significantly higher score for both signature in TGF-CAF with significant downregulation in TGF>>Tie2i-CAF (Figure 6D, S5C-i,ii). This clearly supported the reversal of C2-CAFs towards transcriptional state of C1-CAFs, upon Tie2 inhibition, as observed from pseudo-time analysis.

**Figure 6:**
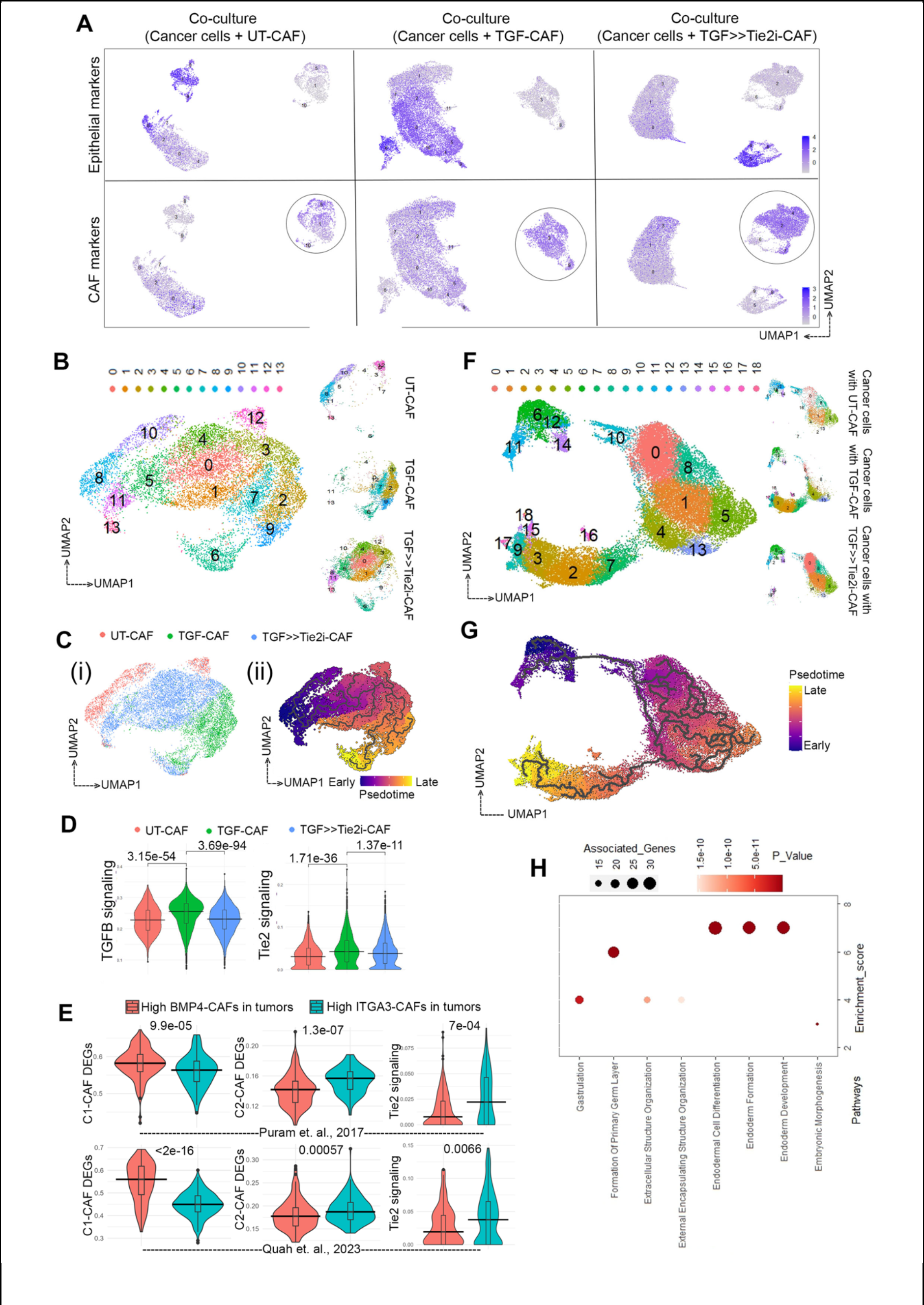
TGFβ-induced myofibroblastic C2-CAFs reprograms oral-cancer cells to acquire embryonic-like transcriptome state. (A) Feature plot showing expressions of epithelial and CAF marker modules on UMAP projection from three conditions of co-cultures as indicated. Circled clusters are annotated as CAF clusters with high CAF marker module scores and negativity for epithelial markers module scores. **(B)** UMAP plot shows re-clustering of CAF clusters from all the three conditions merged, revealing 13 clusters with a total of 11391 cells. A split view of major clusters in a sample specific manner is provided on side panel. **(C)** (i) An UMAP plot visualizing sample wise grouped CAF clusters. (ii) Monocle3 pseudo- time analysis showing CAF dynamic transition along the trajectory. **(D)** Violin plot showing enrichment of TGFβ and Tie2 signaling AUC scores generated by R tool ‘AUCell’, upon TGFβ treatment of CAFs, which was significantly decreased followed by Tie2 inhibition. **(E)** AUC scoring of CAFs from classified patient groups (High BMP4 (C1-like) / High ITGA3 (C2-like)) from Puram et al. and Quah et al. HNSCC datasets shows significant enrichment of C1-CAF DEGs in High BMP4 group, and C2-CAF DEGs and Tie2 signaling in High ITGA3 group. **(F)** Subset of 32354 epithelial cells from all three conditions were merged and re-clustered, identified 16 clusters, projected on UMAP plot. **(G)** Pseudotime analysis exploring transition trajectory of cancer cells. **(H)** Bubble plot showing GO biological process analysis of gene-set among single cell and bulk RNA sequencing of cancer cells co cultured with TGF-CAF. Size of bubble represents numbers of associated genes and colour corresponds to given p value.

To evaluate the presence of C1-type and C2-type CAFs in patient derived samples, we investigated the single cell transcriptome datasets of treatment-naive HNSCC tumors from two independent earlier studies by Puram et. al. and Quah et. al. (39,40). Based on the consistent differential expression of BMP4 and ITGA3 in our current datasets and previously reported study from our group (15), we considered BMP4 and ITGA3 as markers for C1-type and C2-type CAFs, respectively. Patients with higher than median score of BMP4 and lower than the median score for ITGA3 were classified as High-BMP4-CAF patients. Vice versa, individual patients with higher than the median score of ITGA3 and lower than the median score for BMP4 were classified as High-ITGA3-CAF patients (Supplementary Figure S6A, B, C). Crucially, the AUCell scoring was performed for the classified patient groups using scRNAseq- DEGs between our UT-CAF and TGF-CAF (Adj. P value < 0.05) (Table S3). Validating the classification, CAFs in high-BMP4-CAF patient group showed significantly higher score for upregulated UT-CAF-DEGs, whereas; CAFs in high ITGA3-CAFs patient group showed significantly higher score for upregulated TGF-CAF-DEGs (Figure 6E), for both Puram et. al. and Quah et. al. studies. Furthermore crucially in both these studies, CAFs in high-ITGA3 group patients showed significantly higher score for Tie2 signaling, aligning with our observation of Tie2 pathway enrichment in TGF-CAF or C2-CAFs derived from patients (Figure 6E).

### CAF-specific Tie2 activity regulates cancer cell plasticity and stemness in oral cancer cells

Previously we have reported that myofibroblastic C2-CAFs drive stemness in oral cancer cells (15). Therefore, TGFβ induced Tie2-signal in CAFs might act as a potential target against C2-CAFs driven cancer cell reprogramming. To make deeper understanding of the cancer cell reprogramming ability of Tie2-activity in C2-CAFs, we next evaluated the transcriptome state of cancer cells using our co-culture derived scRNAseq data. A total of 32354 cells with higher scores for epithelial gene-set were clustered together from all the conditions to broaden our knowledge on how different subtypes of CAFs modulate cancer cell transcriptome (Figure 6F). Re-clustering patterns of cancer cells revealed 3 major clusters with a total of 18 sub-clusters encompassing different transcriptional states (Figure S7A-i,ii,iii) 6F). While the one major subset of clusters (clusters 6,10-12,14) was common in all three conditions; surprisingly we observed other sets of cancer cells (clusters 0,1,4,5,8,13) shared majorly common clustering neighbourhood when co-cultured with UT-CAF or with TGF>>Tie2i-CAF, suggesting close similarity in their gene expression patterns. Interestingly, a very distinct subset of clusters (clusters 2,3,7,9,15-18) was comprised of cancer cells from TGF-CAF coculture (Table S4), depicting TGF-CAF induced transcriptional reprogramming of cancer cells, which was apparently absent when cancer cells were co-cultured with Tie2- inhibited TGF-CAF (Figure 6F). Pseudotime analysis suggested a dynamics of cancer cell transition trajectory, highlighting that upon co-culture with TGF-CAF this subset of oral cancer cell acquire more evolved state on transition axis with respect to clusters which were unchanged in any co-culture conditions (Figure 6G, S7B). Further, relative position of cancer cells in co-culture with UT-CAF and with TGF>>Tie2i-CAF were almost indistinguishable in axis, implying that Tie2 inhibition may suppress cancer cells reprogramming ability of C2-CAFs.

Emergence of this unique transcriptionally reprogrammed subset of cancer cells upon co-culture with TGF-CAF prompted us to further characterize their molecular nature. We overlapped the differentially upregulated genes in this unique subset of cancer cells (clusters 2,3,7,9,15-18) with genes which were differentially upregulated in cancer cells co-cultured with TGF-CAF in comparison to UT-CAF from our bulk-RNAseq data (Figure 1). 150 DEGs were identified as common among both lists; majorly harboured biological process of early developmental processes, indicating an embryonic-like reprogramming of cancer cells by TGF-CAF (Figure 6H, S7C, Table S5, S6). Taken together, our data clearly suggested that TGFβ induced myofibroblastic C2-CAFs reprograms oral-cancer cells to acquire an undifferentiated phenotype which may have more aggressive functions.

Our bulk-RNAseq analysis revealed a total of 1843 and 1568 genes upregulated and down-regulated respectively in cancer cells co-cultured with TGF-CAF compared to UT-CAF (Table S7). GSEA analysis identified enrichment of signatures for stem cell, EMT, cytokine-cytokine interaction and downregulation of cell-cycle in cancer cells co-cultured with TGF-CAF group (Figure 7A). In support of obtained downregulation of cell cycle marker gene-set; frequency of Ki67-positive cells was found to be reduced in cancer cells co-cultured with TGF- CAF, as compared to UT-CAF (S8A-i, ii). Previous data from our lab has demonstrated an increased frequency of ALDH^High^ stem-like cancer cells (SLCCs) in co-culture with C2-type CAFs (15). Similarly, significantly higher frequency of ALDH^High^ phenotype was observed when oral cancer cells were exposed to TGF-CAF CM (Figure 7B, S8B). Recently we have revealed plasticity in oral cancer cells having ALDH^High^ and ALDH^Low^ phenotype (41). This instigated us to sort ALDH^Low^ cells and coculture with UT-CAF and TGF-CAF for four days. Results clearly suggested that TGF-CAF can significantly favour the shift of ALDH^Low^ cells into ALDH^High^ cells (Figure 7B, S8C). Further, gene expression of cancer cells showed upregulation of stemness associated genes *NANOG*, *OCT4*, *CK14* and *CD44* in two different cancer cell lines exposed to CM of TGF-CAF compared to that of UT-CAF (Figure 7C); clearly suggesting an induction of stemness in cancer cells by TGF-CAF.

**Figure 7.**
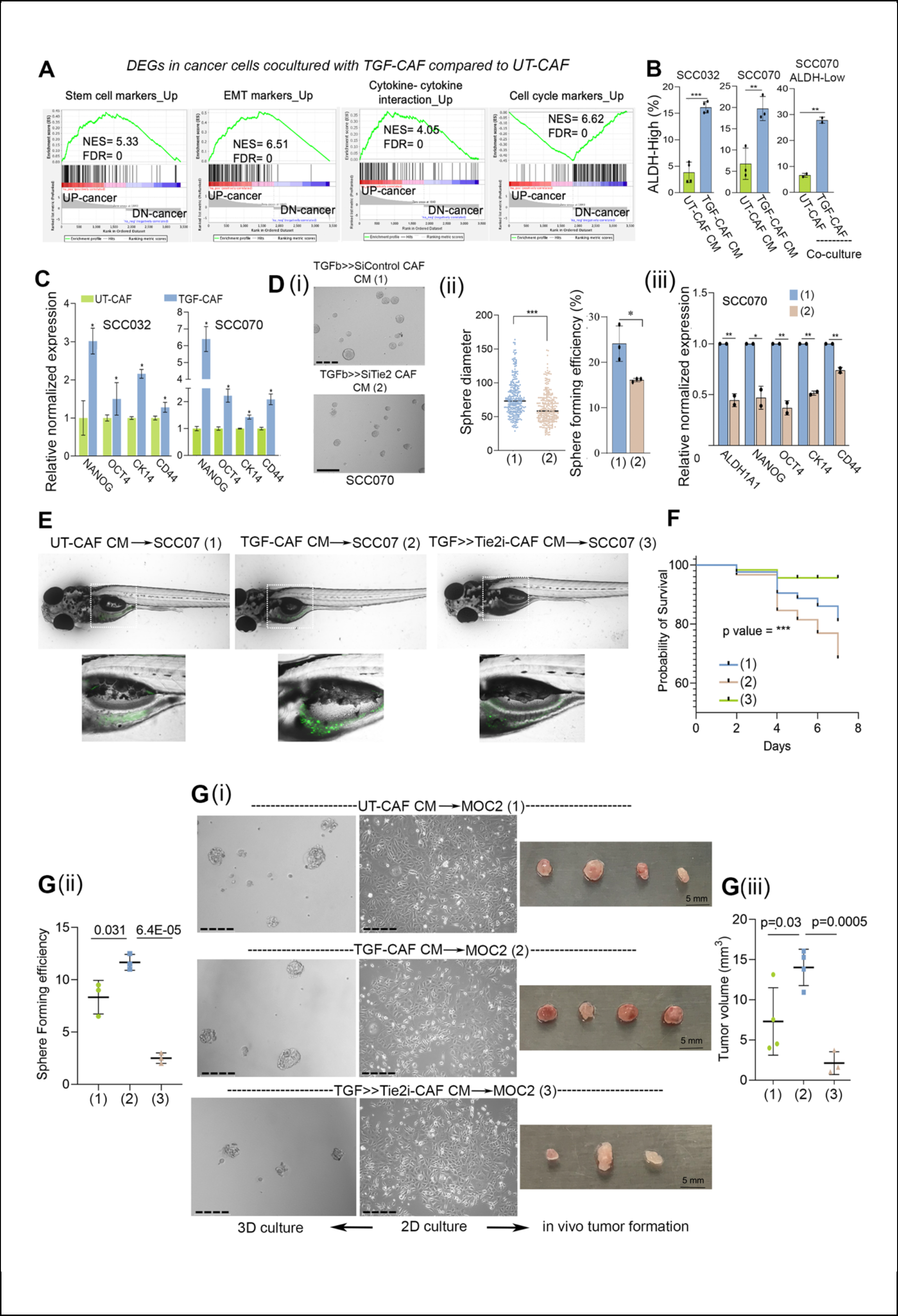
CAF-specific Tie2 regulates cancer cell plasticity and stemness in oral cancer cells. **(A)** Gene set enrichment analysis (GSEA) from transcriptome data of cancer cells, co-cultured with UT-CAF or TGF-CAF for four days. Datasets were obtained from MsigDB database. **(B)** Bar graph showing conversion of ALDH-Low cells into ALDH-High cells upon exposing to conditioned media of UT-CAF or TGF-CAF or upon co-culture as indicated. **(C)** qPCR analysis of stemness associated genes (*OCT4*, *NANOG*, *CD44* and *KRT14* (*CK14*) in two different oral cancer cell lines (SCC070 and SCC032) exposed to CM from KV07 or KV018 CAF, respectively. **(D)** (i) Representative image of 3D-spheroids of SCC070 cell line exposed to CM from TGFβ>>siTie2 or TGFβ>>siControl, followed by testing in spheroid formation assay. (ii) Dot plot showing diameter of formed spheroids of cancer cells from these conditions and bar graph showing sphere forming efficiency of cancer cells exposed to both these conditions. Sphere size was quantified using ImageJ. Spheres of <60 µm diameter were excluded from study. (iii). qPCR analysis of stemness associated genes (*ALDH1A1*, *OCT4*, *NANOG*, *CD44* and *KRT14 (CK14*) in cancer cells following exposed to CM from TGFβ>>siControl or TGFβ>>siTie2 in monolayer culture for 48 hrs. **(E)** Representative images of zebrafish xenografts taken using confocal microscope. GFP positive oral cancer cells (SCC070) were exposed to conditioned media of UT-CAF, TGF-CAF or TGF>>Tie2i-CAF for 48 hrs. Cells were harvested and 100 cells were inoculated into yolk sac of each zebrafish embryo (2-day post fertilization). GFP-positive cell colonies were visible on 4th day of inoculation. **(F)** Kaplan Meier survival plots showing a probability of deaths in zebrafish embryos due to increased tumor burden. **(G)** (i) Representative phase contrast images of MOC2 cells cultured with conditioned media of UT-CAF, TGF-CAF and TGF>>Tie2i CAF for 48 hrs in monolayer culture (2D) and representative images of 3D spheroids of MOC2 cells exposed to CAF-CM from all three conditions as mentioned. (ii) Tree plot showing sphere forming efficiency of MOC2 cells exposed to conditioned media of UT-CAF, TGF-CAF and TGF>>Tie2i CAF. Spheres of <60µ diameter were excluded from study. (iii) MOC2 cells cultured in conditioned media of UT-CAF, TGF-CAF and TGF>>Tie2i CAF for 48 hrs in monolayer culture. These CM exposed MOC2 cells (3x105 cells/mice) were subcutaneously inoculated into syngeneic C57BL/6 mouse models and monitored for 10 days. On day 10th of transplantation, mice were sacrificed and tumors were harvested. Volume of these tumors were measured using ImageJ and plotted in GraphPad prism. **P<0.05, **P<0.01, ***P<0.001. Scale bars, 275 µm.

So far, transcriptome data suggested an ability of TGF-CAF in educating cancer cells to acquire more aggressive transcription state which was significantly suppressed after inhibition of Tie2 activity in CAFs. Therefore, this possibility was next evaluated against stemness in cancer cells by targeting Tie2 expression and activity in TGF-CAF. Oral cancer cell (SCC070) exposed to CM collected from TGF-CAF transfected with Tie2-siRNA (TGF>>siTie2- CAF) showed significant downregulation of tested stemness related genes *NANOG, OCT4, ALDH1A1, CK14* and *CD44* (Figure 7D) as well as spheroid forming efficiency, as compared to CM collected from CAFs transfected with control-siRNA (TGF>>siControl-CAF) (Figure 7D, i-ii). Similar results were obtained with sphere forming efficiency of two different tested oral cancer cell lines. This was significantly increased when exposed to conditioned media of TGF- CAF compared to control (UT-CAF); whereas, it was suppressed when exposed to CM from TGF>>Tie2i-CAF in both the tested cell lines (Figure S9Ai-ii, S9B). Importantly, these same cells were not affected when growing in differentiating adherent condition with serum, suggesting this effect to be specific to stem cell enriching 3D-cultures (Figure S9C). Together, our scRNAseq data analysis and cellular functional assays strongly supported the notion that TGF- CAF-expressed Tie2 plays as one of the most crucial regulators in driving cellular plasticity and maintaining higher stemness in oral cancer cells.

We next evaluated the impact of CAF-induced cancer cell reprogramming on tumor forming ability of oral cancer cells. First, GFP expressing SCC070 oral cancer cell line was exposed to CM obtained from UT-CAF, TGF-CAF or TGF>>Tie2i-CAF for 48 hours. Cells were harvested and 100 cells were injected into yolk sac of each 2dpf (two days post fertilization) embryo. Cancer cell foci formation was monitored under fluorescent microscope for up to seven days and mortality was recorded. Confocal images were taken after 4 days post injection of cancer cells. Interestingly, similar to the results obtained with sphere formation; SCC070 cells incubated with CM of TGF-CAF showed maximum tumor foci formation and also highest mortality of embryos (Figure 7E, F). Interestingly, embryos injected with SCC070 exposed to CM of UT-CAF and TGFβ>>Tie2i-CAF did not show significant cancer cell foci formation with better survival probability, within tested time period (Figure 7E). Importantly, we observed better survival of embryos injected with cancer cells which were exposed to CM of TGFβ>>Tie2i-CAF (Figure 7F). Encouraged from these observations, we next aimed to perform tumor formation assay using murine syngeneic mouse model of oral cancer. Towards this, we first tested if human-CAF-derived CM may exert similar effect on C57BL/6 mouse oral cancer derived cell line, MOC2. Very interestingly, similar to human oral cancer cell lines, sphere forming efficiency of MOC2 cells was significantly increased when exposed to conditioned media of TGF-CAF compared to control and suppressed when exposed to CM from TGF>>Tie2i-CAF (Figure 7G, i-ii). Similar to earlier observation with human cell line, MOC2 cells growing in normal adherent cell culture with serum was not affected, suggesting the effect to be specific to stem cell enriching 3D cultures (Figure 7G-i). Next, these MOC2 cells (3x10^5^ cells/mice) exposed to different CM were allografted subcutaneously into wild- type C57BL/6 mice. Significantly higher tumor volume was observed in conditions where MOC2 cells were exposed to CM of TGF-CAF compared to UT-CAF. In contrast, only 3 out of 4 animals developed tumor and volume of developed tumor was significantly lesser for allografted MOC2 cells exposed to CM of TGFβ>>Tie2i-CAF (Figure 7G-iii). Overall, data clearly supported the impact of Tie2 activity in TGF-CAF, driving cell state transitions of oral cancer cells to acquire stemness.

### Tie2 responsive single cell gene expression data derived modules translate to clinical output of HNSCC patients

Emergence of more aggressive transcriptome state due to the dynamic influence of interaction between the CAF-subtypes and co-cultured cancer cells, prompted us to evaluate the translatability of our observed *in vitro* cellular processes for its clinical significance. The deconvoluted scRNAseq data, where individual cancer cells cocultured with UT-CAF (Red), TGF-CAF (Green) and TGF>>Tie2i-CAF (Blue) were first computed for AUCell scores of EMT and stemness signature) (Figure 8A-i, S10A-i, ii). As anticipated, this pseudo-bulk analysis of scRNAseq data indicated that cancer cells in co-culture with TGF-CAF are enriched with EMT and stemness related genes, and significantly reduced when cancer cells were co-cultured with TGF>>Tie2i-CAF (Figure 8A-ii). Thus, we next mapped these cellular states with the expression signatures of four previously reported molecular subtypes of HNSCC, namely atypical, basal, mesenchymal, classical (Figure 8A-iii) (42,43). While all cancer cells showed very low score for atypical subtype signature; cancer cells cocultured with the UT-CAF (Red) showed significant but marginally higher AUCell score for the classical subtype gene signatures. However, TGF-CAF cocultured cancer cells (Green) had highly significant enrichment of cells with expression pattern for the basal and mesenchymal subtype genes; as shown by AUCell scores (Figure 8A-iv, S10B). Very interestingly, cocultured cancer cells with TGF>>Tie2i-CAF (Blue), retained lower expression of basal and mesenchymal genes signature. Thus, Tie2 activity in TGF-CAFs may drive basal/mesenchymal subtype program in oral cancer cells.

**Figure 8:**
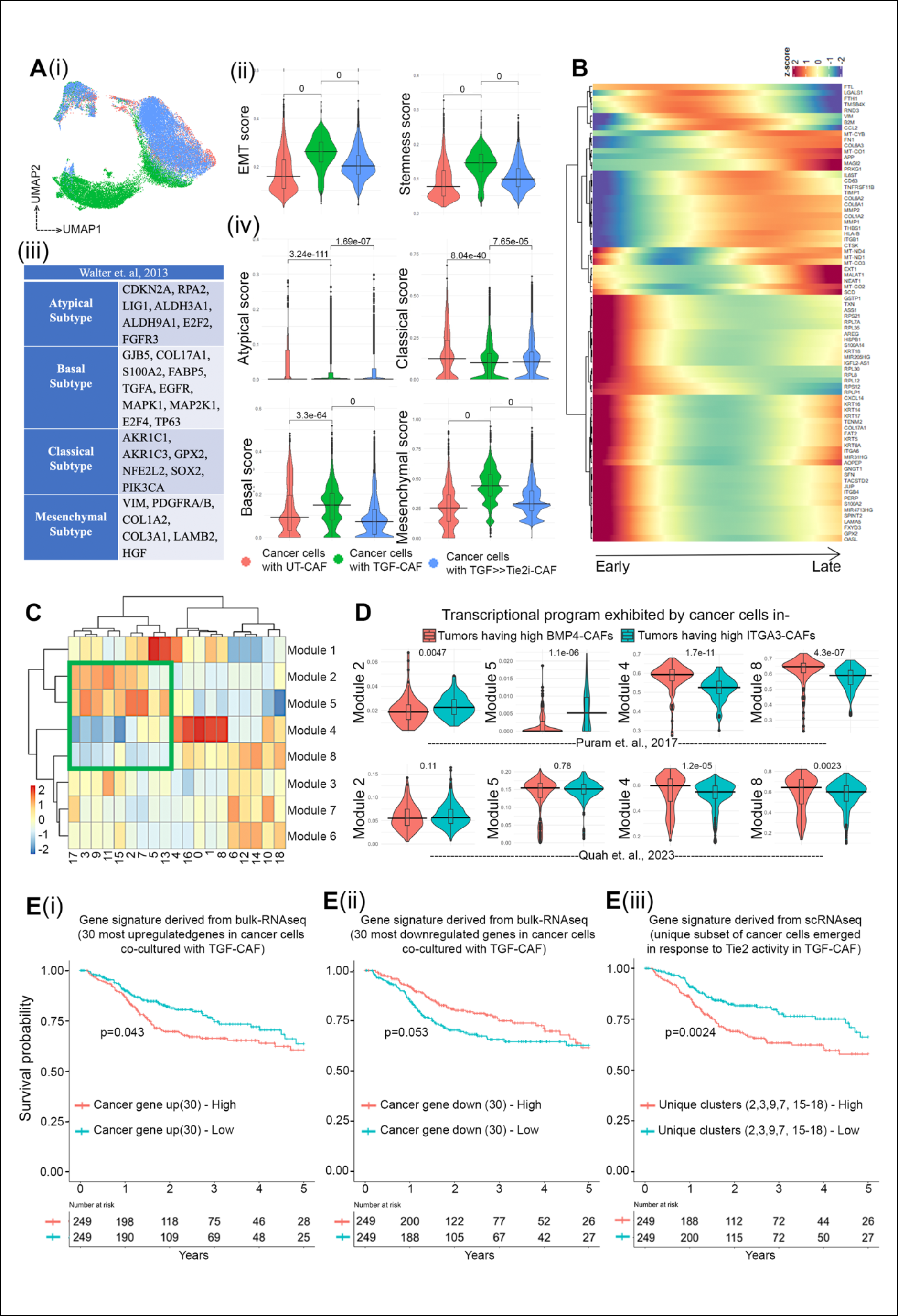
Tie2 responsive single cell gene expression data derived modules translate to clinical output of HNSCC patients. (A) (i) UMAP plot showing colour coded clustering of cancer cells co-cultured with UT-CAF, TGF-CAF and TGF>>Tie2i-CAF. (ii) Violin plot showing EMT and stemness AUC score generated by R tool ‘AUCell’. (iii) Table depicting gene expression based classified molecular subtypes of HNSCC signatures (from walter et al., 2013) (iv) Pseudo-bulk analysis of AUCell scores over cancer cells co cultured with distinct CAF subtypes for the given molecular subtype gene signatures. **(B)** Heatmap showing trajectory variable gene expressions from early to late pseudo-time. **(C)** Expression heatmap of co-regulatory gene modules for each cluster of merged cancer cell subset. Marked green box indicates similar expression pattern of module 2, module 5; and module 4, module 8 on exclusive Tie2 responsive cancer cell clusters. **(D)** AUC scoring of cancer cells from the aforementioned patient groups from Puram et al. and Quah et al. HNSCC datasets shows significant enrichment of modules 4 & 8 in High BMP4 group (C1-like CAF high tumors) in both datasets, and modules 2 & 5 in High ITGA3 group (C2-like CAF high tumors) in Puram et al. and Quah et. al. dataset. **(E)** Prediction of survival probability of TCGA HNSCC patients. Kaplan Meier plot showing survival probability of HNSCC patients harbouring gene signatures of (i)Top 30 upregulated or (ii)Top 30 downregulated genes of TGF-CAF cocultured cancer cells form bulk RNAseq data (iii) Survival probability of patients harbouring gene expression signature obtained from scRNAseq analysis of unique subset of cancer cells co-cultured with TGF-CAF, as mentioned.

To identify the underlying key transcriptional networks as drivers of TGF-CAF-induced cancer cell reprogramming in response to Tie2 activity, we obtained differentially expressed, trajectory variable genes that changed over the pseuodtime shown in figure 6G (Figure 8B). Using this differentially expressed gene set, we constructed coregulatory gene modules of the cancer cells which resulted in eight dynamically regulated gene modules across all single cell clusters of cancer cells. Very interestingly, module 2 and 5 were collectively upregulated and module 4 and 8 were downregulated in cluster 2,3,5,7,9,11,13,15, and 17 (Figure 8C). Interestingly, except cluster 5 and 11 all these clusters were mainly contributed by subsets of oral cancer cells in response to Tie2 activity in TGF-CAF. The downregulated modules (4 and 8) showed regulation of translational process and upregulated module (2 and 5) showed the process of cell junction organization and cell migration (Figure S10C).

We next examined if these modules are operated in cancer cells *in situ* in presence of C1-type and C2-type CAFs within the oral tumors. For this, we utilized our previously classified patient groups (Figure S6A,B,C) and evaluated the single cell gene expression pattern of malignant cell population from primary tumor of two independent studies by Puram et. al. and Quah et. al. (39,40) (supplementary Figure S10 D,E). Dimensional reduction of malignant cells subsets from the individual patients in High-BMP-CAF or High-ITGA3-CAF group showed marked difference in the gene expression patterns among these patient groups in UMAP projections (Supplementary Figure S10E). AUCell scoring was performed for this classified patients groups using uniquely expressed genes in modules 2, 5, 4 and 8 (Suppl. Table S8). To our excitement, we observed that the malignant cells from high-ITGA3-CAF patient group showed significantly lower score for modules 4 and 8 for both Puram et. al. and Quah et. al. studies and higher score for modules 2 and 5 in Puram et al study (Figure 8D). Thus, this analysis has provided evidence of *in situ* reprogramming of cancer cell by CAF-specific Tie2- activity within HNSCC and therefore may have its clinical translatability.

To provide clinical interpretation of observed biology the prognostic significance of the data was next evaluated. We used DEGs between cancer cells co-cultured with TGF-CAF in comparison to UT-CAF from our bulk-RNAseq data and correlated with expression data of HNSCC patient cohort in TCGA study. Survival analysis was performed using gene-set specific ssGSEA score. Patients with their individual ssGSEA scores, more than mean were classified as ’high’, and others as ’low’. Survival of these groups was estimated using Kaplan–Meier (KM) curves and Cox-regression analyses. Interestingly, among all comparisons (Figure S11A,B); patients with higher ssGSEA-score for top 30 upregulated genes showed poorer 5-year disease specific survival (Figure 8E-i); whereas, top 30 downregulated genes showed better survival (Figure 8E-ii). Our scRNAseq data has discovered emergence of specific subsets of oral cancer cells with more evolved transcriptome state in response to Tie2 activity in TGF- CAF. Therefore, we next tested the gene-set of this unique subset of cancer cells, as alternate signature. Patients with higher ssGSEA-score for this signature also showed significantly poorer 5-year disease specific survival (Figure 8E-iii, S11C); highlighting clinical relevance of observed CAF-specific TGFβ-ANGPT-Tie2 signaling axis driven reprogramming of oral cancer cells.

## Discussion

Studies on Angiopoietin/Tie2 pathway have been majorly focused on endothelial cell functions, related to angiogenesis and vessel maturation (25,44). However, our work has identified the role of TGFβ-signaling in epigenetic downregulation of *ANGPT2*, leading to Tie2- activation in oral-CAFs, with TGFβ-ANGPT-Tie2 to be regulating each other in a closed loop. Since, depletion of endogenous TGFβ or Tie2 in primary C2-CAFs or TGF-CAF significantly upregulated the levels of *ANGPT2* expression with concomitant decrease in Tie2- phosphorylation and myofibroblast phenotype of CAFs, we suggest that TGFβ-induced *ANGPT2* downregulation may be the key event in induction of Tie2 signaling and maintenance of C2-CAF state. As one of the possible mechanisms of *ANGPT2* downregulation, we identified the role of histone deacetylases (HDACs) in TGFβ-induced H3k27-deacetylation of the *ANGPT2* promoter, where *ANGPT2* and other tested C1-CAF associated genes showed significantly reduced suppression in presence of inhibitor of class-I HDACs, valproic acid; suggesting HDAC-regulated induction of myofibroblastic CAF-state. Similarly, TGFβ mediated HDAC7 repression of *PPARGC1A* gene was also found to be crucial for fibroblasts activation in fibrotic lung tissue (45). Further work will be needed to dissect the precise mechanism behind ANGPT-Tie2 axis in maintaining TGFβ-induced myofibroblast-phenotype of CAFs. In endothelial cells, Tie2 signaling activates small GTPase Rac1 through PI3K and Akt, leading to its localization on adherence junction (46). However, the mechanisms by which ANGPT-Tie2 signal impacts formation of focal adhesions, cytoskeleton remodelling and stress-fiber arrangement is still under exploration in endothelial cells (47).

CAFs as major co-existing component of complex tumor ecosystem, exhibit dynamic molecular interactions to cooperate and co-evolve in tumor microenvironment (48,49). Several studies have correlated high abundance of stromal myofibroblastic, αSMA-positive CAFs with poor prognosis of oral cancer patients (12,50–52); however, studies exploring CAF- driven mechanisms have been limited in oral cancer. Our previous report had demonstrated the role of myofibroblastic C2-CAFs in providing more conducive microenvironment for enhanced stemness (15). Advancing our understanding, the current study identified CAF- specific Tie2-signaling in TGFβ-induced CAFs, reprogramming malignant cells to embryonic cell-like state; suggesting as one of the mechanisms generating stemness-supporting niche in oral tumor microenvironment. Since, CM was sufficient for educating cancer cells and cytokine-cytokine receptor interaction was one of the most significant gene-sets enriched in co-cultured cancer cells; we suggest that secretory factors from Tie2-activated CAFs may drive cancer cell reprogramming to acquire stemness in oral tumor. Although, studies have suggested the role of TGFβ-induced CAFs in supporting tumorigenic ability of cancer cells (53,54); however, further work will be required to demonstrate the specific Tie2-mediated factors secreted from TGF-CAF in driving oral cancer progression.

TGF-CAF showed myofibroblastic phenotype with certain overlapping similarities with CAF-types reported earlier in OSCC tumors. Activation of CXCL9/10/11-CXCR3 axis is shown recently in TDO2^+ve^ myofibroblasts present in OSCCs (55). Similarly, a recent study performed with T1-stage OSCC tissue, with matched dysplasia and adjacent normal tissue reported a subcluster of CAFs as mesen_CAFs showing certain resemblance with TGF-CAF; like, enrichment of TGFβ, EMT, angiogenesis, and PI3K-AKT-mTOR pathways (56) or earlier defined classical myofibroblast marker αSMA (39). Significantly, our defined CAF-subtypes specific gene signatures as well as Tie2-pathway signature was detected in fibroblast clusters in two independent single cell studies of HNSCC tumors (39,40). Thus, the Tie2-induced cellular processes exhibited by TGF-CAF in our study showed resemblance with the subpopulation of CAFs within oral tumor microenvironment of HNSCC patients. This highlighted the possibility of CAFs to undertake endothelial-like transition and our model system may be appropriate for studying the biology of such transitions in future.

Few clinical trials are being attempted to directly target stromal CAFs in solid tumors. Although targeting TGFβ is successful in pre-clinical models, it faced major problems when tested under clinal trials, owing to its dual role (57–59). Reversal of pro-tumorigenic state by reprogramming CAFs using vitamin A and D has been demonstrated (60). Since, Tie2-active CAFs reprogram oral cancer cells to acquire aggressive phenotype; CAF-specific function of Tie2 may provide therapeutic benefit. Supporting the possibility, ‘Rebastinib’ as a novel and selective inhibitor of Tie2 is currently under clinical trials against leukaemia and locally advanced and metastatic solid tumors in combination with chemotherapy (61,62).

Our single cell transcriptome data facilitated us in profiling dynamic changes influenced by interactions between the CAF-subtypes and co-cultured cancer cells. Cancer cells showed enrichment of the signature of mesenchymal- and basal-subtype of oral cancer after being reprogrammed by TGF-CAFs. Similar malignant-basal subtype specific gene expression was previously found to be positively associated with partial-EMT phenotype and negatively associated with differentiation state in malignant cells (39). In connection to this, The Tie-2 activity in TGF-CAF was found to facilitate co-cultured oral cancer cells to acquire embryonic-like cell state with increased stemness and EMT signatures. Similarly, earlier studies with enrichment of embryonic stem cell signature correlated with aggressive cancer behaviour and poorer prognosis of oral cancer patients (63–65). Moreover, CAF- reprogrammed cancer cell specific gene expression modules was also present in malignant cell population in these HNSCC patients. Intriguingly, specific genes which were differentially expressed in subset of cancer cells in response to CAF-specific Tie-2 activity acted as prognostic signature in HNSCC patient cohort. Therefore, the CAF-specific Tie2-activity may act as a causal link between the abundance of myofibroblastic C2-CAFs in tumor stroma and associated poor prognosis. Overall, our study has opened an avenue of targeting Tie2- signaling in CAFs to combat against aggressive behaviour of oral cancer.

## MATERIALS AND METHODS

### Cell lines and culture conditions

The human oral cancer cell line SCC070 and SCC032 were obtained from Dr. Susanne M. Gollin, University of Pittsburgh (Pennsylvania). SCC070 cells were maintained in Dulbecco’s Modified Eagle Medium (DMEM/F12) medium and SCC032 cells were maintained in Minimal Essential Medium (MEM) and supplemented with NEAA, glutamine and antibiotics and 10% fetal bovine serum. GBC02 and MOC2 were cultured in Epilife and IMDM/F12 media respectively, supplemented with FBS and growth factors. Oral Cancer associated fibroblasts (CAFs) were grown in DMEM/F12 medium supplemented with 10% FBS, 1X ITS and antibiotics. Cells were cultured in an adherent monolayer condition at 37°C under a humidified atmosphere containing 5% CO2. Human subjects were included in this study (EC/GOVT/01/12) after approval obtained from the institutional ethics committee of National Institute of Biomedical Genomics (NIBMG) and the institutional review board of Tata Medical Center (TMC), Kolkata, India.

### Conditioned media generations

TGFꞵ induction (10 ng/ml) was done in low serum media (DMEM/F12 supplemented with 1% serum and 0.1X ITS) for 48 hrs. Tie2 inhibitor (500 ng) was added either before one hour of TGFꞵ treatment (pre-treatment) or following TGFꞵ induction (post-treatment) for 48 hrs. Cells were washed with serum free media and conditioned media was collected after another 48 hrs.

### Collagen-I-matrigel contraction assay

TGFꞵ induction (10 ng/ml) was done in low serum media (DMEM/F12 supplemented with 1% serum and 0.1X ITS) for 48 hrs. Cells were harvested and mixed with collagen I-matrigel and for another 48 hrs. A matrix of rat tail collagen I (354249, corning) with final concentration of 4.6 mg/ml and matrigel (354234, corning) with final concentration of 2.2 mg/ml was prepared. 1N sterile NAOH (S2770, Sigma) was used to neutralize acetic acid of collagen I-matrigel matrix and diluted in PBS. The gel plug was detached from the sides to allow contraction. Gel plug with no cells was used as control. Matrix contraction was monitored and imaging was done by Chemidoc imaging system (BioRad).

### siRNA mediated knockdown

CAFs were transfected with 50nM of siRNA of Tie2, ACTA2, ANGPT1 and ANGPT2 and scrambled siRNA as a negative control (Eurogentec) for 48 hrs in low serum media using INTERFERin kit (Cat# 409-10; Polypus) following manufacturer’s instructions.

### Lentivirus transduction

The PLenti.CMV.GFP.Puro (Addgene#17488) plasmid were transduced into the SCC070 cells. Recombinant Lentivectors were produced by transient transfection of transducing vectors into HEK293T cells with packaging vectors, a plasmid expressing the HIV-1 gag/pol, tat, and rev genes. Lentiviral vector mediated transduction was validated by GFP reporter expression in fluorescence microscopy and FACS based phenotyping was done to get the pure population of the transduced cells, GFP positive cells were then sorted and expanded.

### RNA extraction, cDNA preparation and RT-PCR

Adherent cells were washed twice with ice-cold PBS and immediately lysed in RLT-plus buffer (Cat. # 74104, Qiagen) containing β-mercaptoethanol. Total RNA was extracted using RNA- Easy plus kit (Cat#74034, Qiagen) as per the manufacturer’s instructions. Total RNA was measured in the Nanodrop quantification system (Thermo-Scientific). 500 ng of total RNA was converted to cDNA using iScript advanced cDNA synthesis kit for RT-qPCR (Cat# 1725037, Biorad) according to the instructions of the manufacturer.

The expression level of stemness-related genes was performed in CFX96 real time qPCR machine (Biorad) using iTaq™ Universal SYBR® Green Supermix (Cat# 1725121). The primers were designed using Primer Blast (NCBI). Housekeeping gene GAPDH was used as internal control. The expression levels of mRNAs were normalized to GAPDH (dCT) and the relative normalized expression of test sample was calculated as the fold change relative to the control (2^^-^ddCT).

### ALDEFLUOR assay

The ALDH activity of oral cancer cells co-cultured with CAF was measured using the ALDEFLUOR assay kit (Cat. #01700, StemCell Technology), according to the manufacturer’s protocol. DEAB (Diethylaminobenzaldehyde) was used as a negative control for ALDH assay. Anti-human CD90-BV421 antibody (1:50, cat # 562556; BD) was used to stain the CAF and propidium iodide was used to identify non-viable cells. CD90 negative cancer cells were checked for ALDH staining and analysis was done using the FACS-Aria Fusion instrument (Becton Dickinson). Frequency of ALDH high cells were analyzed using FCS express software.

### Sphere formation assay

Cancer cells were seeded with DMEM/F12 media, supplemented with 10% FBS. After 24 hours cells were washed with plain media and conditioned media was added for 48 hrs. After 48 hrs, cells were harvested. Single cell suspension of oral cancer cells was made in serum-free medium (DMEM/F12) supplemented with 20 ng/ml epidermal growth factor (Cat #PHG0311,Sigma), 20 ng/ml bFGF (Cat #PHG0261, Invitrogen), hydrocortisone and B27 (cat #12587010) and mixed with geltrex (Cat #A14132-02). 300 viable cells per well was seeded in ultra-low adherent 96-well plates (Corning, Cat#3474). The growth factors were supplemented on alternate days. 3D spheres were monitored and imaged on the seventh day and quantified using ImageJ software.

### Bulk RNA sequencing and pathway analysis

In brief, CAFs and cancer cells were co-cultured and following co-culture, both the cell types were separated by FACS. RNA extraction was done from a pure population of individual cell types and bulk RNA sequencing was done using Illumina HiSeq2500. Quality assessment was done by using Agilent Bio-analyser 2100 with Agilent nano kit. The library was prepared by TruSeq Stranded Total RNA Library prep kit. The data was processed by the core facility of National Institute of Biomedical Genomics, India. Significant (p value=<0.05) differentially upregulated and downregulated (log2 fold change>= 2) genes were explored for pathway enrichment analysis by Gene Set Enrichment Analysis (GSEA) and Cytoscape software.

### Zebrafish xenograft

Zebrafish were maintained at a 28°C incubator as approved by the NIBMG committee. 48 hrs post fertilized embryos were dechorionated and 100 GFP positive cells were injected in the yolk sac cavity. Post injection, embryos were maintained at 32°C incubator. Fish were monitored and deaths were recorded up to seven days post injection and survival analysis was done using GraphPad prism. Fluorescent imaging was performed using EVOS M7000 and Nikon laser scanning confocal microscopy after anesthetizing using tricaine solution.

### Mouse allograft

Five-week-old C57/BL6 mice were subcutaneously inoculated with 3*10^5^ MOC2 cells exposed to CAF conditioned media. Mice were housed in sterile cages, maintained in a temperature-controlled room and were fed autoclaved food and water. Ten days post cell inoculation, tumors were harvested by sacrificing the mice. Volume of the tumors were measured using ImageJ software and graphs were generated using GraphPad prism. All animal experiments were done as approved by the animal welfare committee of the IISER Kolkata and NIBMG- Kalyani (IISERK/IAEC/2020/014).

### Immunofluorescence

CAFs were plated on poly-D-Lysine and Collagen-I coated 100 mm tissue culture dishes. Cells were fixed with 4% paraformaldehyde at room temperature and washed with ice cold PBS. Permeabilization and blocking was done with 0.2% Tween 20 and 10% goat serum in PBS respectively. Cells were incubated with primary antibodies targeting aSMA (ab7817, 1:300), Ki67 (ab16667, 1:100), Tie2 (A7222,1:100), pTie2 (AF2720, 1:100), BMP4 (ab39973, 1:150) overnight at 4 ⁰C. Cells were incubated with alexa Fluor conjugated goat anti-mouse and goat anti-rabbit secondary antibodies for one hour at room temperature at a dilution of 1 μg/mL. Nucleus was counter-stained with DAPI (ProLong® Diamond Antifade Mountant, Cat. # P36962 Thermo). Image acquisition was done by EVOS M7000 (ThermoFisher) and Nikon confocal laser scanning microscope (Ti2, Nikon, Japan). Cells were quantified using ImageJ software.

### Quantification of total and phosphorylated protein

For total protein quantification, intensity of each cell was measured in a gray scale format and normalized with the mean area of the cells. The formula used for quantifying total protein was “CTCF = Integrated Density – (Area of selected cell X Mean fluorescence of background readings)”. Phospho-Tie2 puncta was quantified using ImageJ software. Images were converted in grayscale and length and width of each bright pTie2-puncta was measured. Puncta which were more than 2µ in length were considered as mature phosphorylation marks, anything below was regarded as immature phosphorylation and discarded from the analysis.

### Immunohistochemistry

Surgically dissected oral tumor tissue samples were obtained from Tata Medical Hospital, Kolkata. These sections were used for IHC analysis of aSMA and Tie2 protein expression.

Tissue block was rehydrated, and antigen retrieval was done in the sodium citrate buffer (pH- 6). The samples were then incubated with 3% H2O2 for 10 min. Tissue sections were incubated with monoclonal aSMA (1:50) and polyclonal Tie2 (1:50) antibody overnight at 4°C. Tissue sections were incubated with HRP-conjugated secondary antibody and stained with DAB (Vector, Cat# SK-4105), dehydrated and mounted with DPX (Vector, Cat# H-5000). All the pictures were obtained using the EVOS M7000 microscope (ThermoFisher). Stromal aSMA and Tie2 expressions were scored on a scale of 1 to 4 (10%; 10%–25%; 25%–50%, and 50% positive).

### Chromatin Immunoprecipitation (ChIP) pull down and ddPCR

4.3 million cells of C1 CAF with and without TGF-β were cross-linked using 1% formaldehyde for 10 min at 37°C, and then 0.125 M of glycine was added to quench the remaining. After thorough washing with ice-cold PBS, the cell pellets were resuspended in RIPA, and sonicated using covaris (M200) in glass microtubes (microTUBE AFA Fibre Pre-Slit Snap Cap 6X16mm) to obtain DNA fragments between 200 to 1,000-bp. The lysate was then divided into six equal fractions and 5% of it was retained for Input control. Target proteins were precipitated using antibodies against HDAC2 (ab124974), P300 (ab14984), H3 (ab176842), Lys-27 trimethylated histone H3 (ab6002) and Lys-27 acetylated histone H3 (ab177178) (all purchased from Abcam, MA, USA). Normal IgG was used as negative control. Antibody and lysate were mixed by rotating overnight in 4°C at 13 rpm. Protein A/G-magnetic beads washed with 0.01% PBST followed by ChIP dilution buffer. It was then added to each condition and left them to rotate at 13 rpm for 4 h at 4°C to collect the immune-precipitated complexes. After washing, the beads twice with each buffer i.e. ChIP low salt buffer, high salt buffer, LiCl Buffer and TE Buffer, the DNA was eluted using the elution buffer and kept overnight at 65°C, 650 rpm for de-crosslinking. The following day, elutes were treated with proteinase K for 3h at 56°C, 650 rpm. DNA was extracted by the phenol/chloroform method and ethanol precipitation and re- suspended in 50 *μ*l water for CHIP samples and 200 *μ*l for input-control. ddPCR was carried out to check for the presence of histones at C1-signature gene, BMP4 (between region -708bp and -628bp) & ANGPT2 (-360bp & -1600bp) region.

The ChIP pulldown DNA and Input-control DNA was used to perform digital droplet PCR. Evagreen Qx200 ddPCR specific reagents were used. The regions selected were TATA binding and Initiator regions for ANGPT2 while in case of BMP4 no specific TATA binding site could be located hence the region near Initiator was chosen to ensure presence of active and suppressing chromatin marks with relatable expression of genes. The entire ChIP results represented were after normalization using IgG value followed by percentage calculation against H3, the positive control for all the experimental pulldowns.

### Single cell RNA sequencing library preparation and sequencing

Co-cultured cells were harvested and made a single cell suspension by incubating with accutase in 37°C for 3-4 minutes followed by filtering through a 70 μm cell strainer. After a brief centrifugation at 300g for 3 minutes, cell pellets were resuspended in the cultured low serum media. Using trypan blue, the cells were counted and checked for viability. The suspension having more than 80% cell viability was diluted to a concentration of approximately 1000 cells/µL. Chromium controller (10X Genomics), chromium 3ʹ library, Gel bead kit and finally the chromium Single cell 3ʹ chip kit (10X Genomics) were used according to the manufacturer’s instruction to make Single cell gel bead-in emulsion and for sequencing library construction. Agilent high sensitivity chips in TapeStation were used to examine the quality of the libraries and the quantification was done using QuantStudio-7 real time PCR. All the libraries were pooled and sequenced as a paired-end manner in Illumina NovaSeq-6000. Sequence depth was kept at an average of 50,000 raw reads per cell and per sample 200 million paired-end reads were kept as a standard measure. Base call files were converted into FASTQ and GRCh38 reference transcriptome was utilized for the read alignment using CellRanger 4.0.0 (10X Genomics). R package Seurat 4.3.0 was used for the quality control (QC) and the subsequent analysis from the generated gene-barcode matrix from CellRanger.

### QC, clustering, and cell type annotation

Cells having less than 10% mitochondrial read and total genes expressing in between 400 and 8000 were taken for further analysis. Genes that were expressed in less than 3 cells were discarded. Seurat function ‘LogNormalize’ was used to normalize the expression for each cell by considering the feature count of each cell divided by the total count for that cell multiplied by the default scaling factor set by Seurat i.e., 10000. Variance stabilizing transformation (VST) method was implied to identify the most variable genes i.e., 2000 genes for our case, for principal component analysis (PCA). We constructed graph-based clusters using 15 principal components (PCs) with a resolution of 0.75 and visualized by Uniform Manifold Approximation and Projection (UMAP) using Seurat function ‘RunUMAP’. The Seurat object metadata automatically then stores cluster identities and the sample information in seurat@ident and slots, respectively. Probable doublets were removed as described previously (Jerby-Arnon et al., 2018) and again clustering was carried out. For cell type annotation we assigned a module score of established gene sets of epithelial and CAF markers (Puram et al., 2017) for each cell by ‘AddModuleScore’ function which considers the average expression of the gene set in all the cells from all the clusters. Cell types were segregated by higher or lower scores for the module scores. CAF clusters were separated based on higher CAF gene set module scores and lower or negative for epithelial signature scores. The rest of the clusters were taken into account as epithelial cell clusters. We merged the cell type specific clusters by ‘merge’ function and normalized, scaling, clustering was done again as described earlier.

### Identification of DEGs of clusters and gene set enrichment analysis

DEGs were identified using Seurat function ‘FindAllMarkers’ with co-parameters such as only.pos=F, and logfc.threshold = 0.25 and the enrichment analysis was performed at cluster level. We used ‘Enrichr’ for gene set enrichment and gene ontology analysis. KEGG and Molecular Signature Database (MSigDB) for hallmark gene sets, were used as reference databases for all the pathway enrichment analysis. For all the gene set enrichment analysis, we used free publicly available GSEA software (http://www.gseamsigdb.org/gsea/msigdb/human/annotate.jsp). For computing signature scores, gene signatures were taken either from Puram et al., 2017, cell or from MsigDB gene sets. Significant pathways (FDR<0.05) were taken for further analysis.

### Pseudotime analysis

Trajectory inference was carried out by monocle3 for the cell type specific merged Seurat objects. SeuratWrappers was used to convert the Seurat object to a cell data set object by the function ‘as.cell_data_set’ to provide the input object for monocle3. The cluster information and the other data calculated in Seurat metadata was copied in the cell data set object for getting started with trajectory building. Pseudotime value for each cell was then stored internally by learn_graph method and starting root node was provided to assign the start of the trajectory by ordering the cells. Differentially expressed genes along the trajectory were then identified by Morans’s I test method built in monocle3. Co-expressed or co-modulated genes were clustered into modules by find_gene_modules function and cluster wise gene module expression were analyzed.

### Gene signature scoring / AUC scoring

Gene signature scores were generated by R tool ‘AUCell’ which uses area under the curve (AUC) measurement to construct the extent to which a critical subset of our input gene signatures were expressed within all the expressed genes in all the cells. A rank for all the genes based on the degree of expression were built for each cell by ‘AUCell_buildRankings’ function and then with ‘AUCell_calcAUC’ function we configured a matrix of scores for the specific gene set on each cell of all the clusters. The median value of AUC scores for all the clusters of CAF/epithelial cell type was used for our analysis.

### Single sample gene set enrichment (ssGSEA) and survival analysis

The differentially expressed genes (DEGs) in the treated case were obtained using DESeq2 (Love et al., 2014). The DEGs were shortlisted by adjusted p-value < 0.05 and ranked by their log2-fold change. The top 30 of the upregulated and downregulated genes for the bulk samples were selected as the gene sets for further analysis. In contrast, all the upregulated genes were considered for the single-cell sample, as there were no downregulated genes in the shortlist. These gene sets were used to score the TCGA-HNSC samples (The Cancer Genome Atlas Network, 2015) using the GSEApy (Fang et al., 2023) implementation of the single sample gene set enrichment analysis (ssGSEA) algorithm (Barbie et al., 2009). This method evaluates the enrichment of the genes in the gene set compared to the background expression and provides a score quantifying the activity of the genes. Patients were stratified into high-scoring and low-scoring groups based on the median ssGSEA scores, and Kaplan-Meier (KM) analysis was done to compare the 5-year Disease-specific survival (DSS).

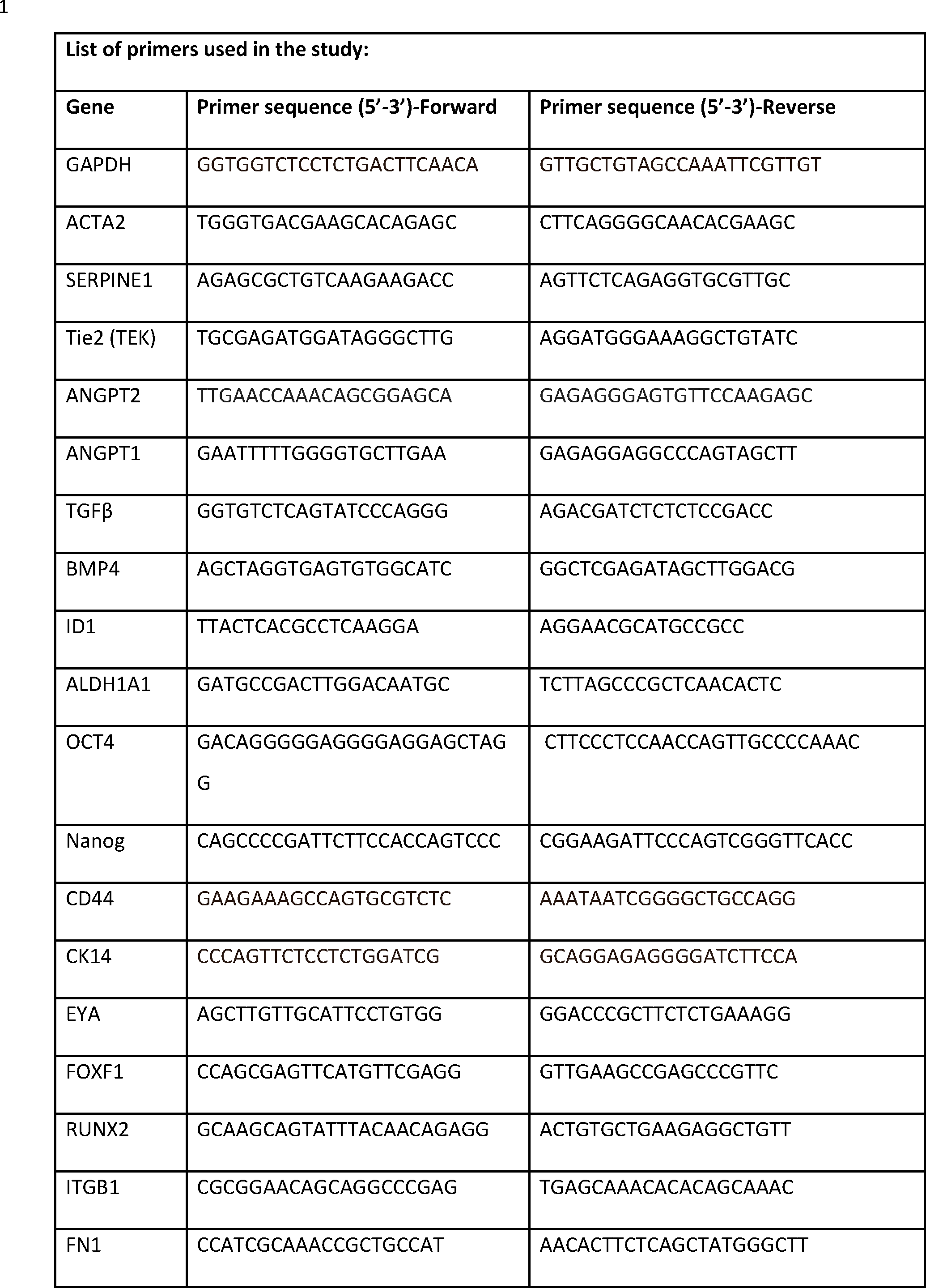
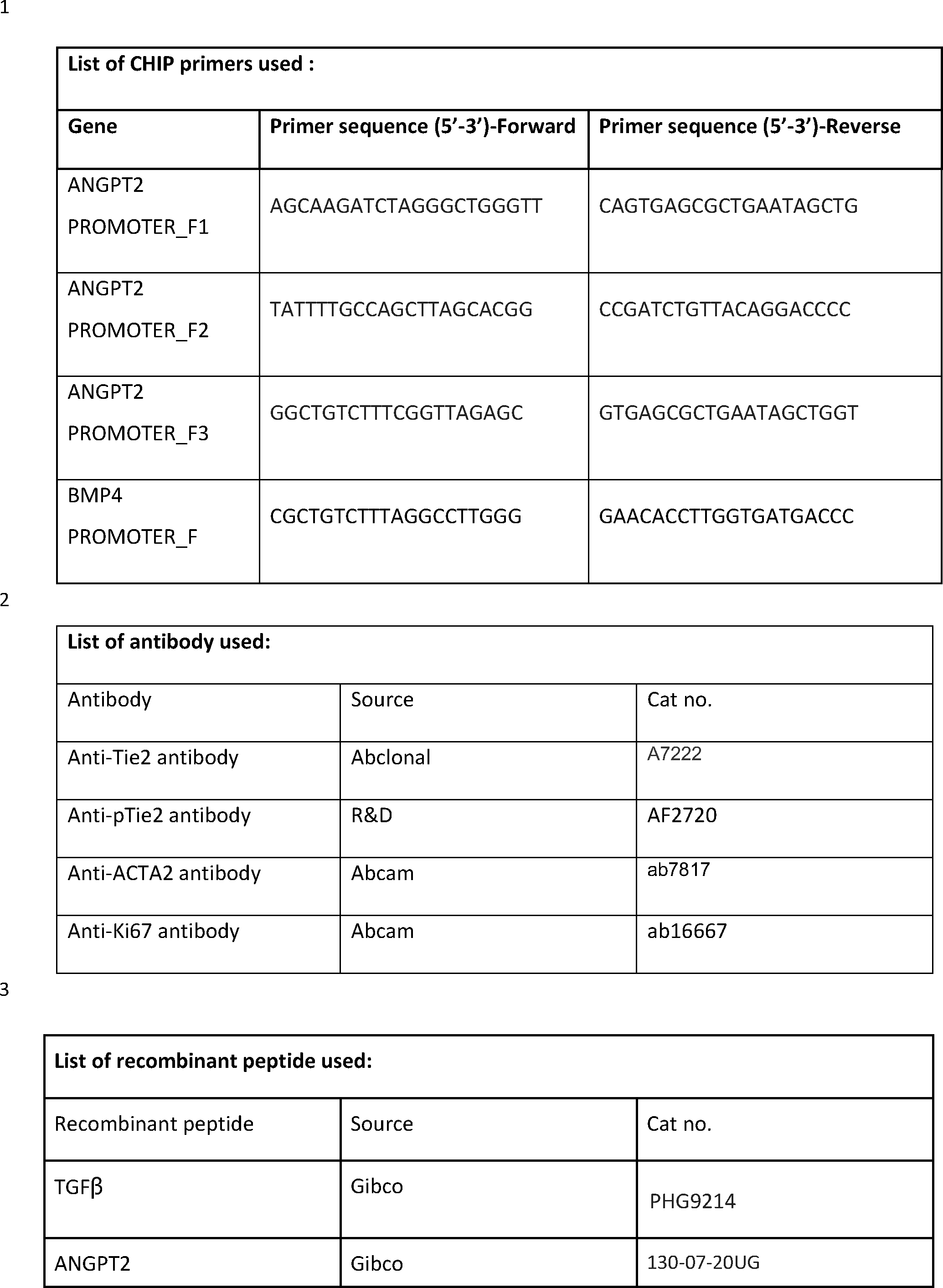
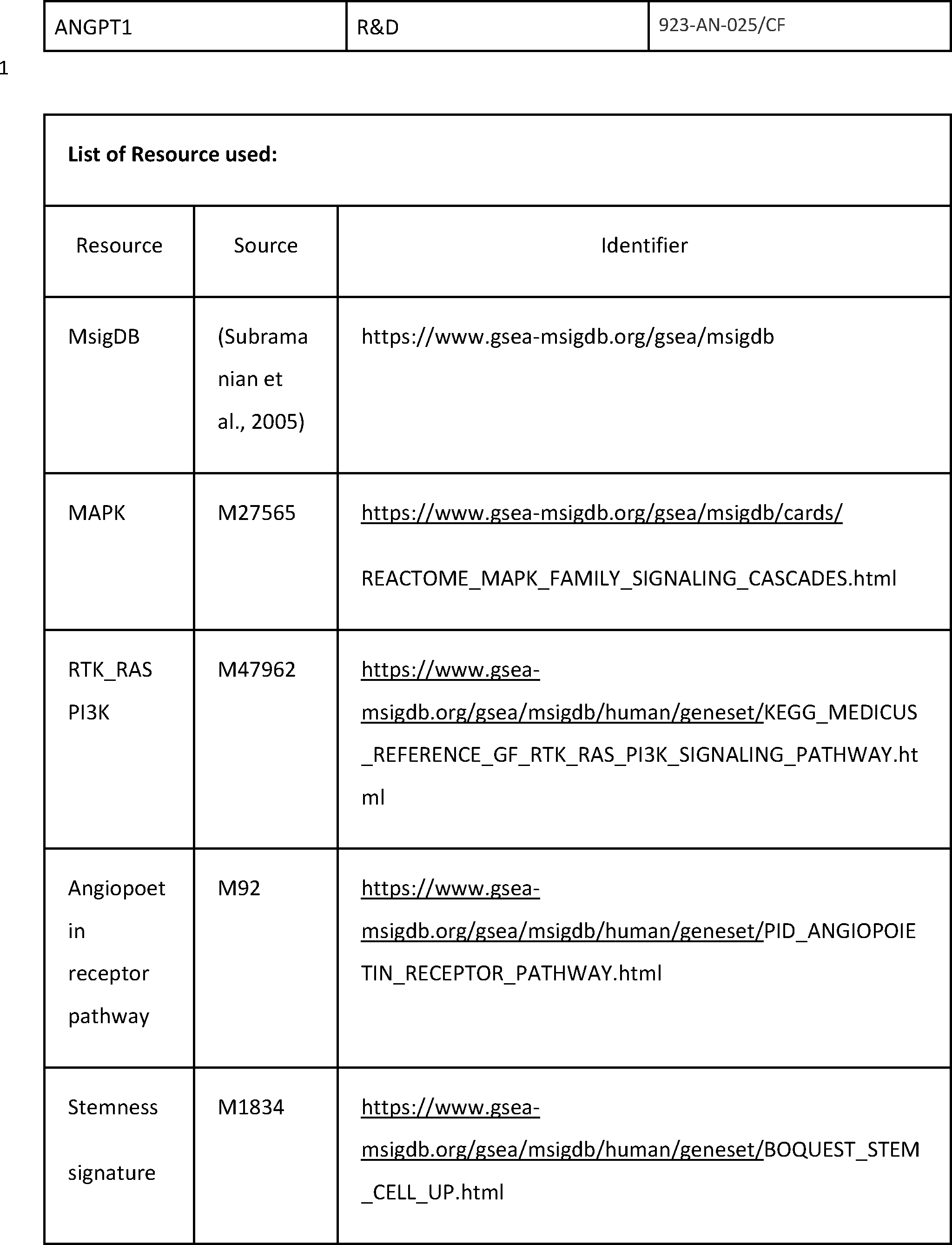
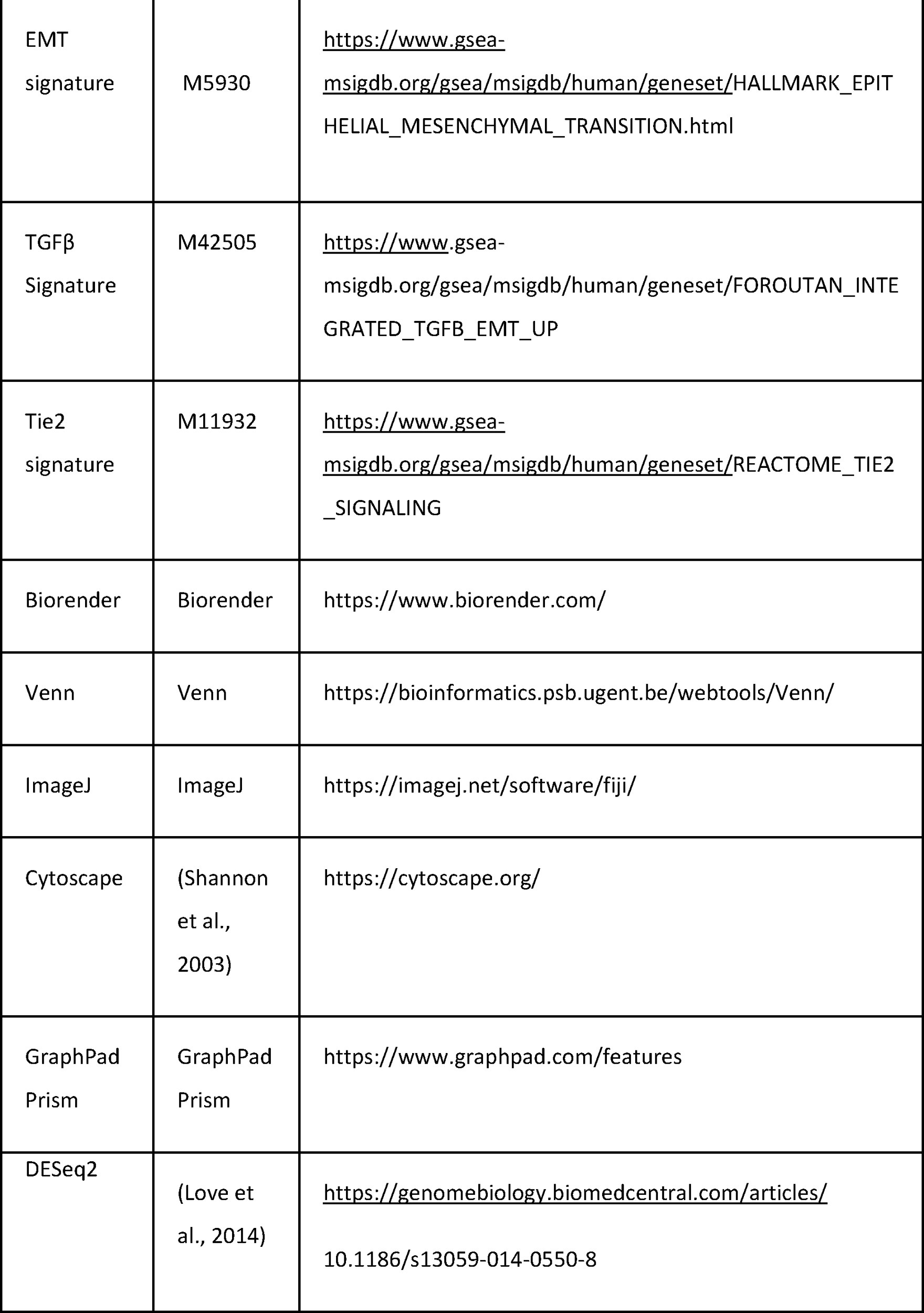
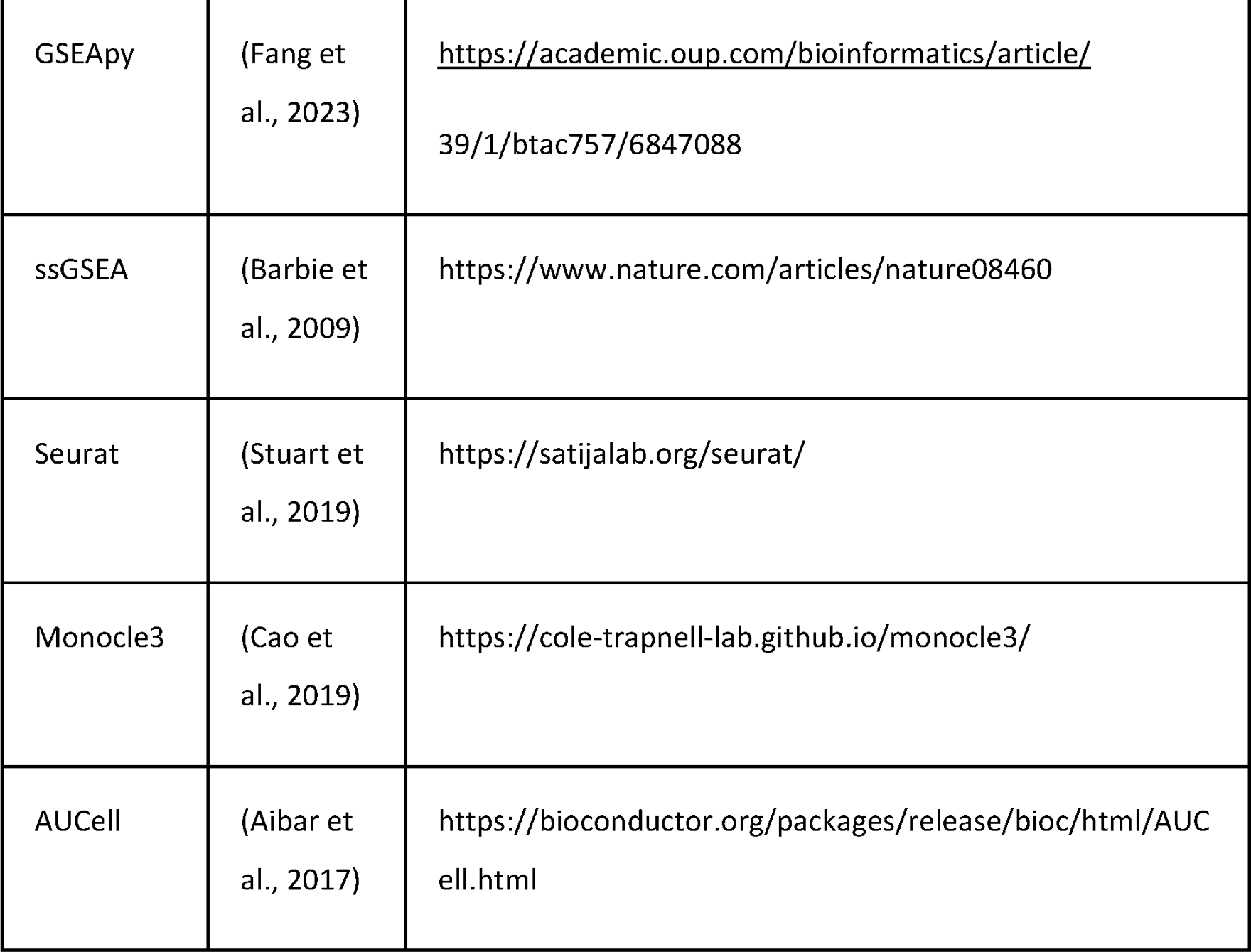

## Contributions

Conceptualization- S.S., P.M.; Methodology- P.M., U.S., P.P., A.K.P., K.J.S., B.V.H., A.G., S.S.R., S.R., and S.S.; Investigation- P.M., U.S., K.J.S., P.P., A.K.P., S.S.; Formal Analysis- P.M., U.S., K.J.S., P.P., B.V.H., S.R., A.G., N.K.B, A.M., and M.K.J.; Writing Original Draft- P.M. U.S., K.J.S., P.P. and S.S.; Review and Editing- P.M., U.S., K.J.S., P.P., A.K.P., S.K.M., B.V.H., M.K.J. A.G., N.K.B., S.S.R., M.A. S.R., A.M. and S.S.; Resources- S.S., N.K.B., M.K.J., A.M., M.A., J.D.S., S.K.M., R.S., P.A.; Supervision-S.S.; Funding Acquisition-S.S

## Funding

This work is supported by the intramural funds from NIBMG to SS.

### Acknowledgement

P.M. and P.P. acknowledges CSIR, India, U.S. acknowledges UGC, India, H.B.V acknowledges PMRF, India for fellowship support. M.K.J. acknowledges the support of philanthropic donation from Param Hansa Philanthropies. Syngeneic mouse experiment was performed in Animal facility of IISER-Kolkata supported by cluster project of department of biotechnology, Govt. of India. We acknowledge Prof. Kartiki V. Desai and Dr. Malancha Ta for suggestions and advice.

---

**S1:**
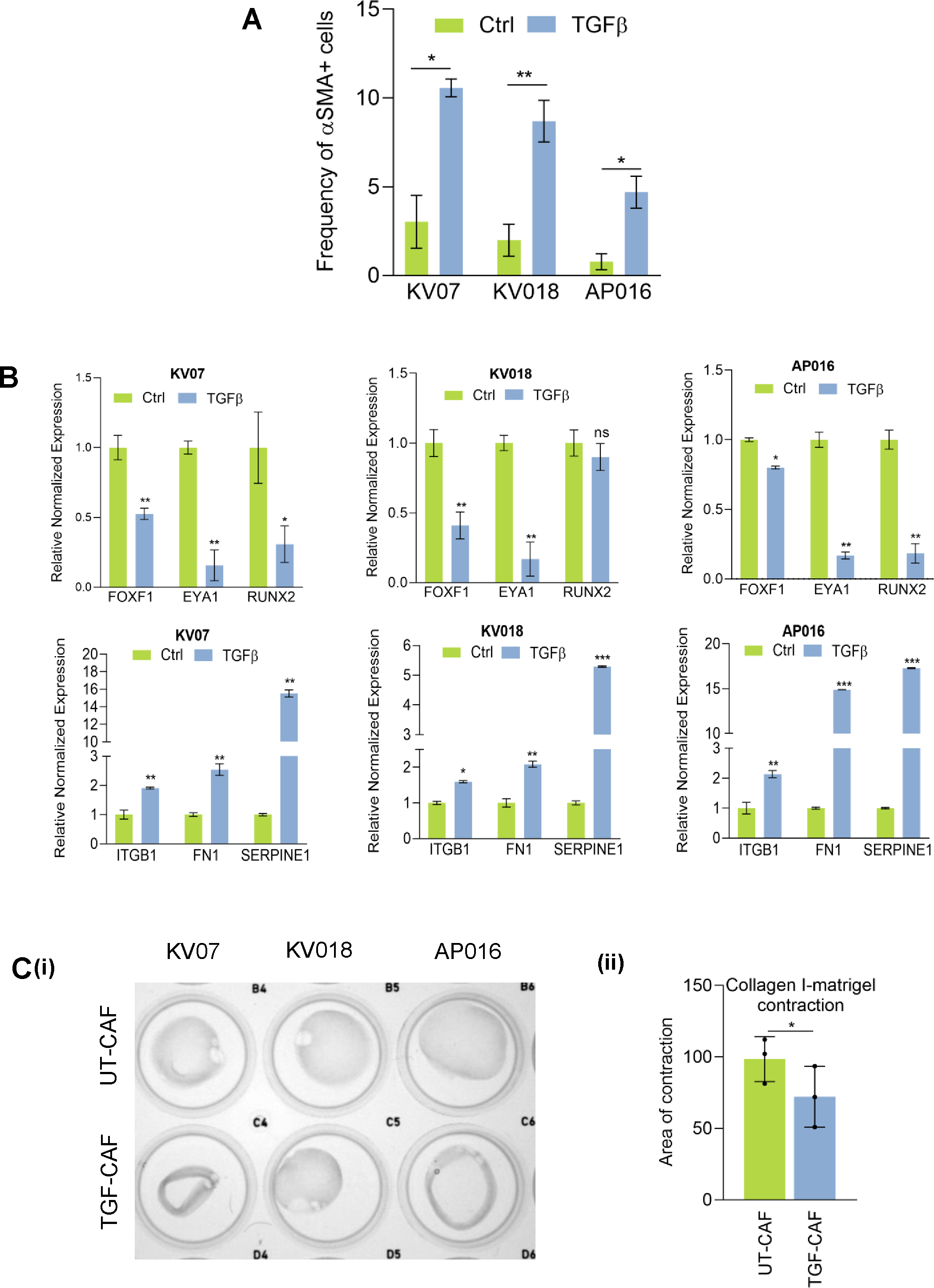
A. Quantification of myofibroblasts frequency following 10 ng/ml TGFꞵ treatment for 48 hrs. B. qPCR analysis of C1 CAF (*EYA1*, *RUNX2*, *FOXF1*) and C2 CAF (*SERPINE1*, *ITGB1*, *FN1*) markers in CAFs stimulated with 10 ng/ml TGFꞵ for 48 hrs. C. (i) Image showing ‘Collagen I-matrigel’ matrix-plug contraction ability of UT-CAF and TGF-CAF. (ii) Result shows Mean± SD of matrix area contraction of three biological repeats ( three different CAFs). Paired student T test was calculated. *P<0.05, **P<0.01, ***P<0.001 .

**S2:**
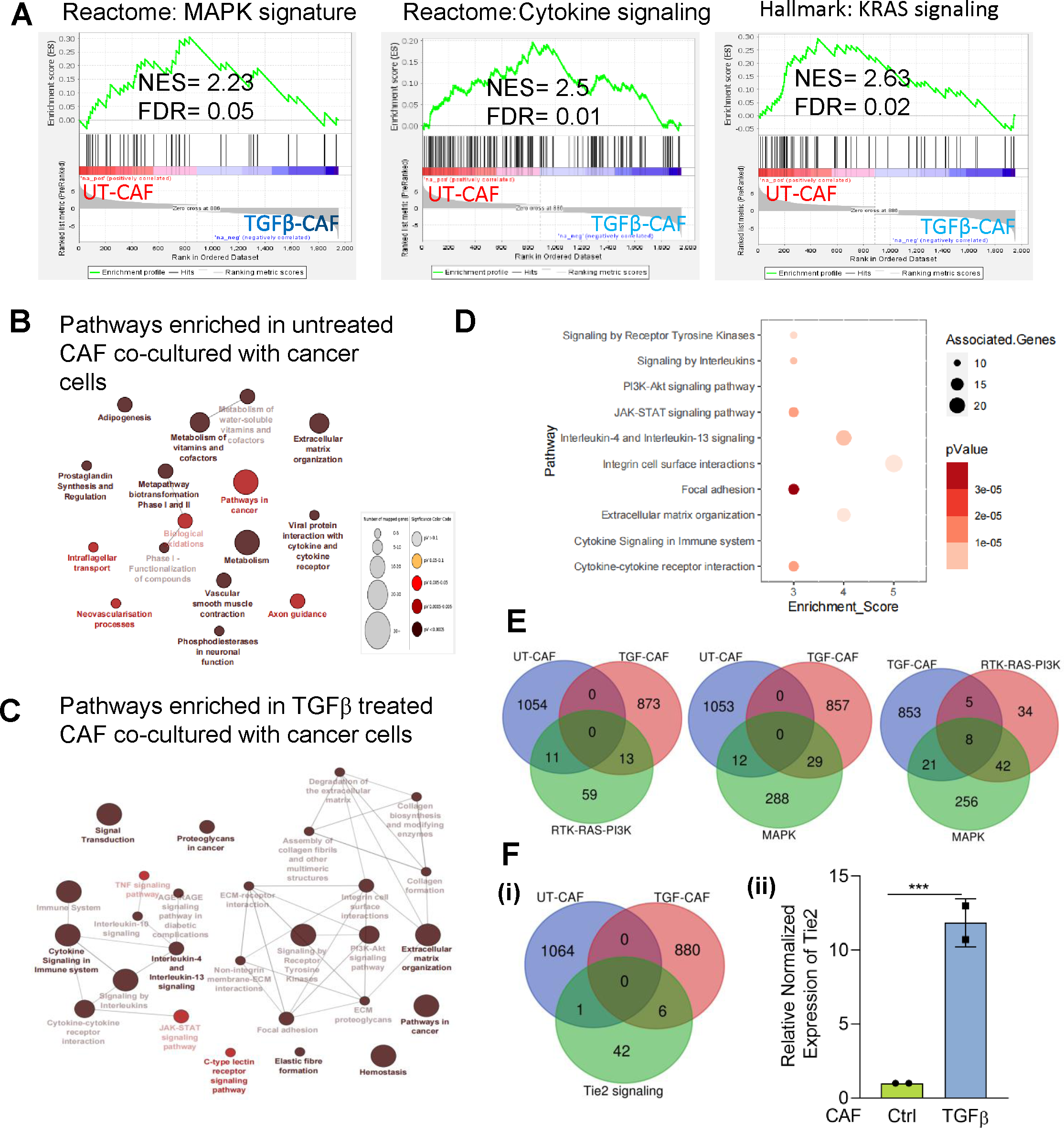
A. Gene set enrichment analysis (GSEA) of CAF stimulated with 10 ng/ml and co-cultured with cancer cells for 4 days. MAP kinase pathway and cytokine-cytokine interaction were positively enriched. Red and blue lines indicated that genes upregulated and downregulated in TGF-CAF co-cultured with cancer cells. Datasets were obtained from MsigDB database. B-C. Cytoscape analysis of positively and negatively enriched networks in TGF-CAF co-cultured with cancer cells. Color coding represents p value and size of nodes represents the number of mapped genes in a particular pathway. D. Pathways shown were found significantly enriched (p value < 0.001) in the TGF-CAF co-cultured with cancer cells. The TGF-CAF signature includes 886 upregulated genes defined with LogFC > 1 and adjusted p value < 0.05 in compared to UT-CAF co-cultured with oral cancer cells. E. Venn diagram showing overlapping gene expression between RTK-RAS-PI3K, MAPK pathway and TGF-CAF co-cultured with cancer cells for 4 days. F. (i) Venn diagram showing overlapping gene expression between Tie2 pathway pathway and TGF-CAF co-cultured with cancer cells for 4 days. (ii) qRT-PCR showing abundance of Tie2 mRNA in TGF- CAF co-cultured with cancer compared to the UT-CAF co-cultured with cancer cells. F. *P<0.05, **P<0.01, ***P<0.001 .

**S3:**
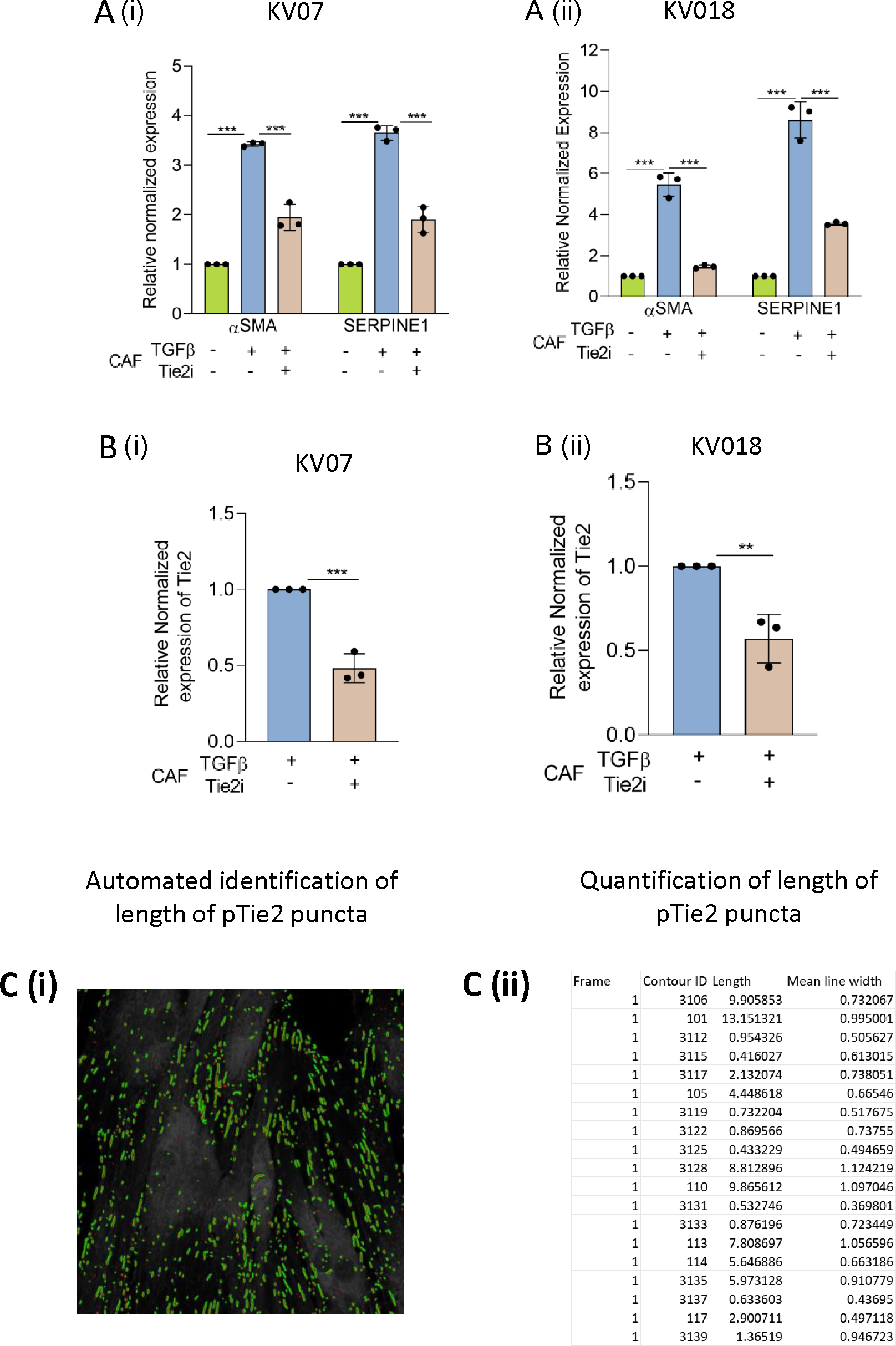
A(i, ii) qPCR analysis of C2 CAF markers (*αSMA* and *SERPINE1*) following Tie2 inhibition in TGF-CAF of KV07 and KV018 respectively. B(i, ii) Tie2 inhibitor was added for 48 hrs after TGFβ induction (TGF-CAF>>Tie2i) and qPCR for *Tie2* was performed. C(i) Representative images revealing automated quantification of pTie2 puncta. Puncta’s with high contrast were measured. (ii) Puncta’s length <2 µm were considered immature puncta and discarded from analysis. Puncta’s length >2 µm were mature puncta and quantified. Paired student t-test was performed between tested samples; *P<0.05, **P<0.01, ***P<0.001 .

**S4:**
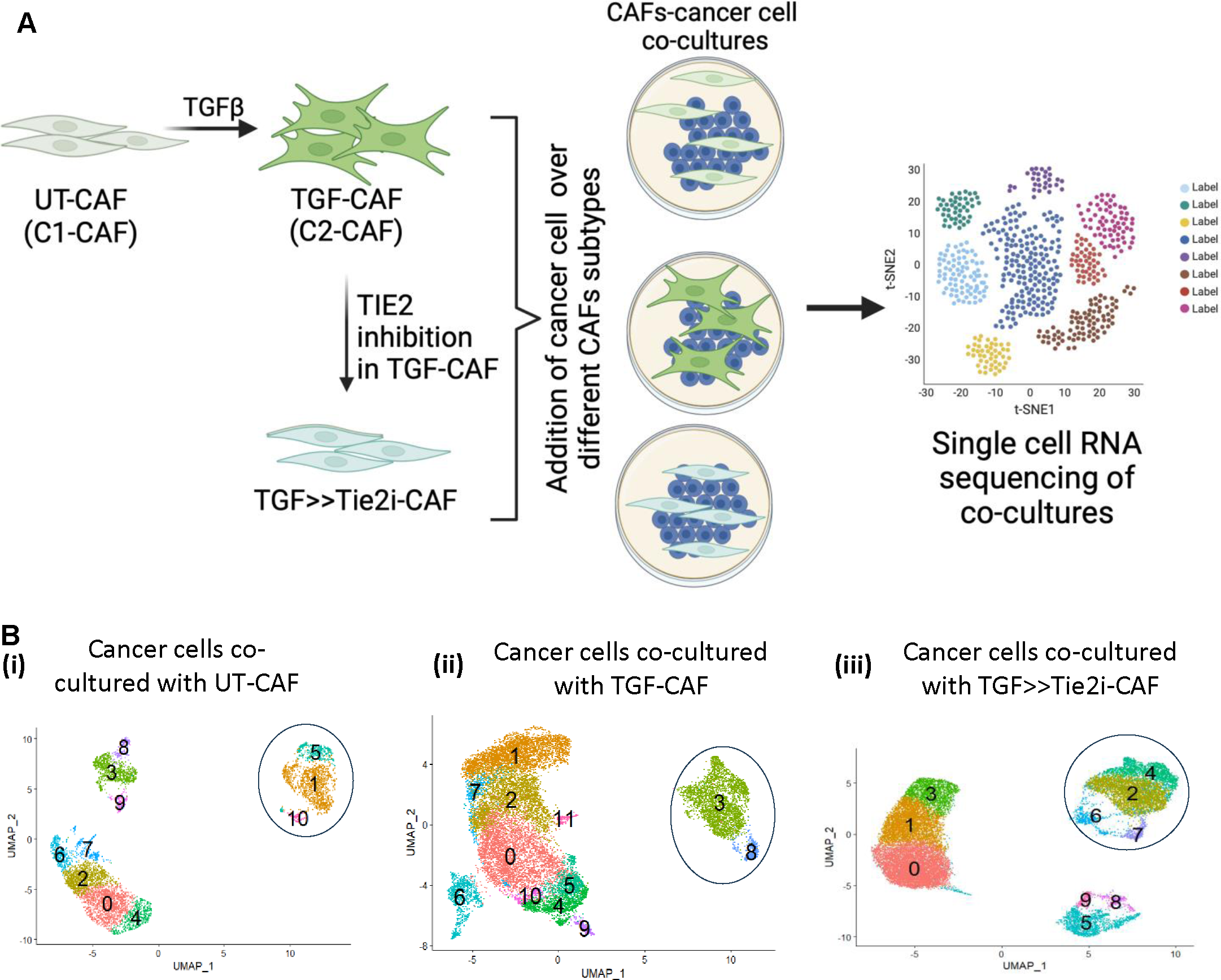
A. Schematic for scRNAseq experiment. B. Clusters of single cells from CAF-cancer cell coculture on UMAP projection. Encircled clusters indicate CAF clusters.

**S5:**
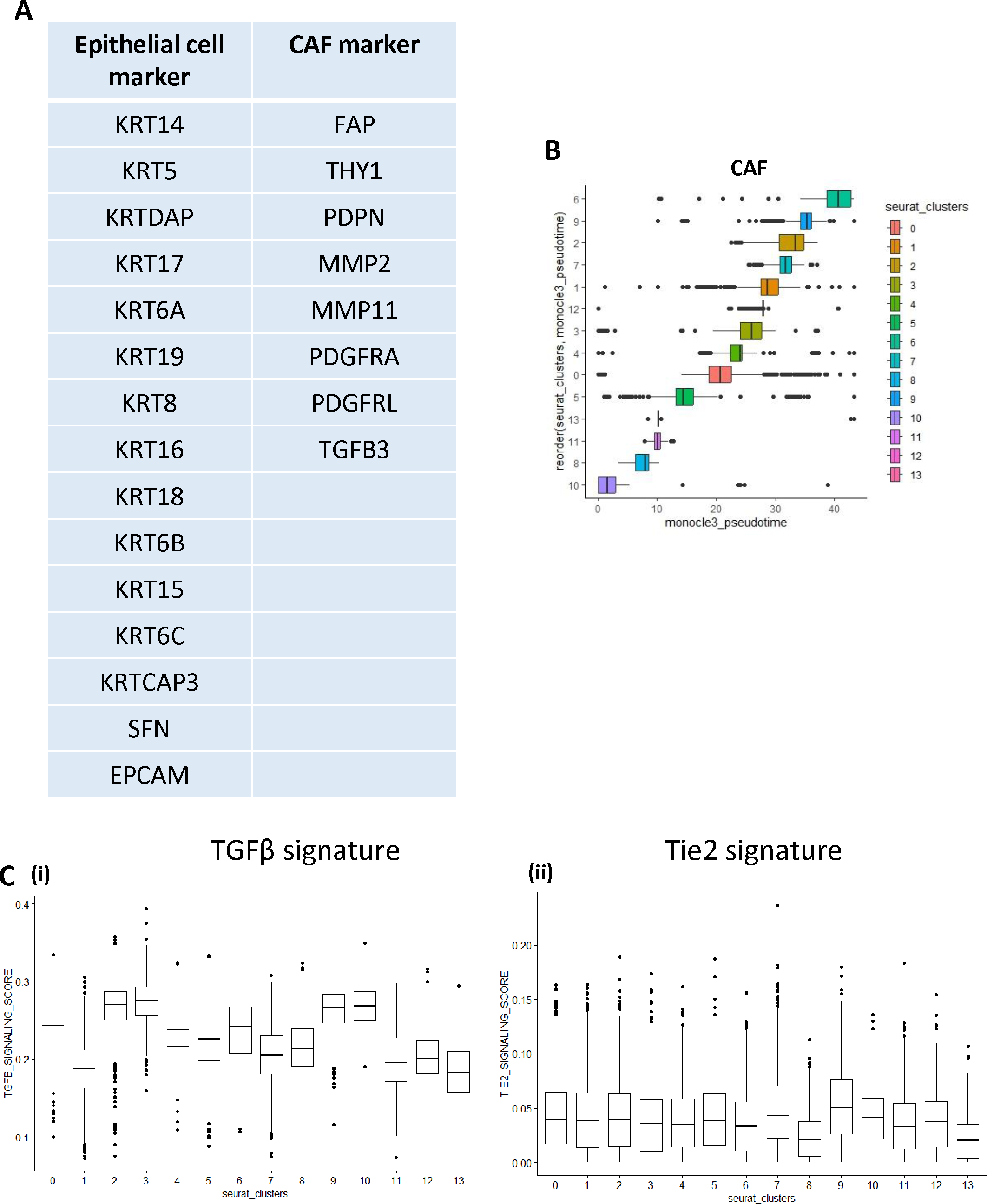
A. List of canonical markers to identify and annotate epithelial and CAF cells in co-cultured scRNA seq clusters. B. Boxplot showing distribution of Monocle3 Pseudotime for each cluster of merged CAF conditions. C. Box plot showing the distribution of TGFꞵ and Tie2 signaling signature AUC scores across different clusters calculated for each single cell. AUC score measurement was performed by the R tool ‘AUCell’. TGFꞵ and Tie2 signaling signature was obtained from Molecular Signature Database (MsigDb)

**S6:**
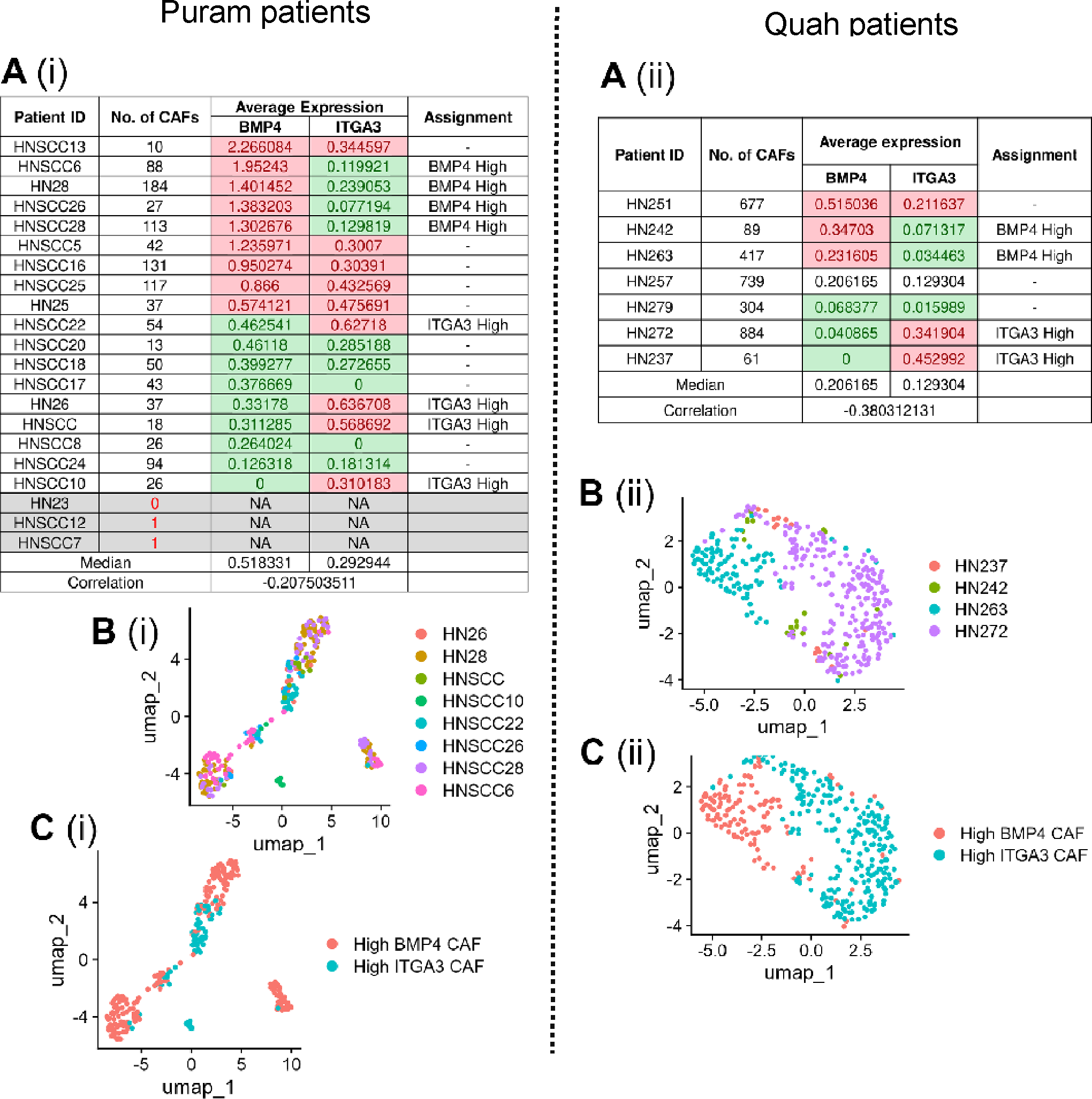
A. (i) Classification of patients from Puram et al. and (ii) Quah et al.as BMP4-High/ ITGA3-High groups with the criterion being above-median expression for one marker and below-median expression for the other. Patients with < 3 identified CAFs and unclassified patients were excluded from further analysis; B. (i - ii) UMAP plots of CAFs from the classified patients, colored based on patient-wise sample-origin, or C. (i-ii) based on assigned grouping.

**S7:**
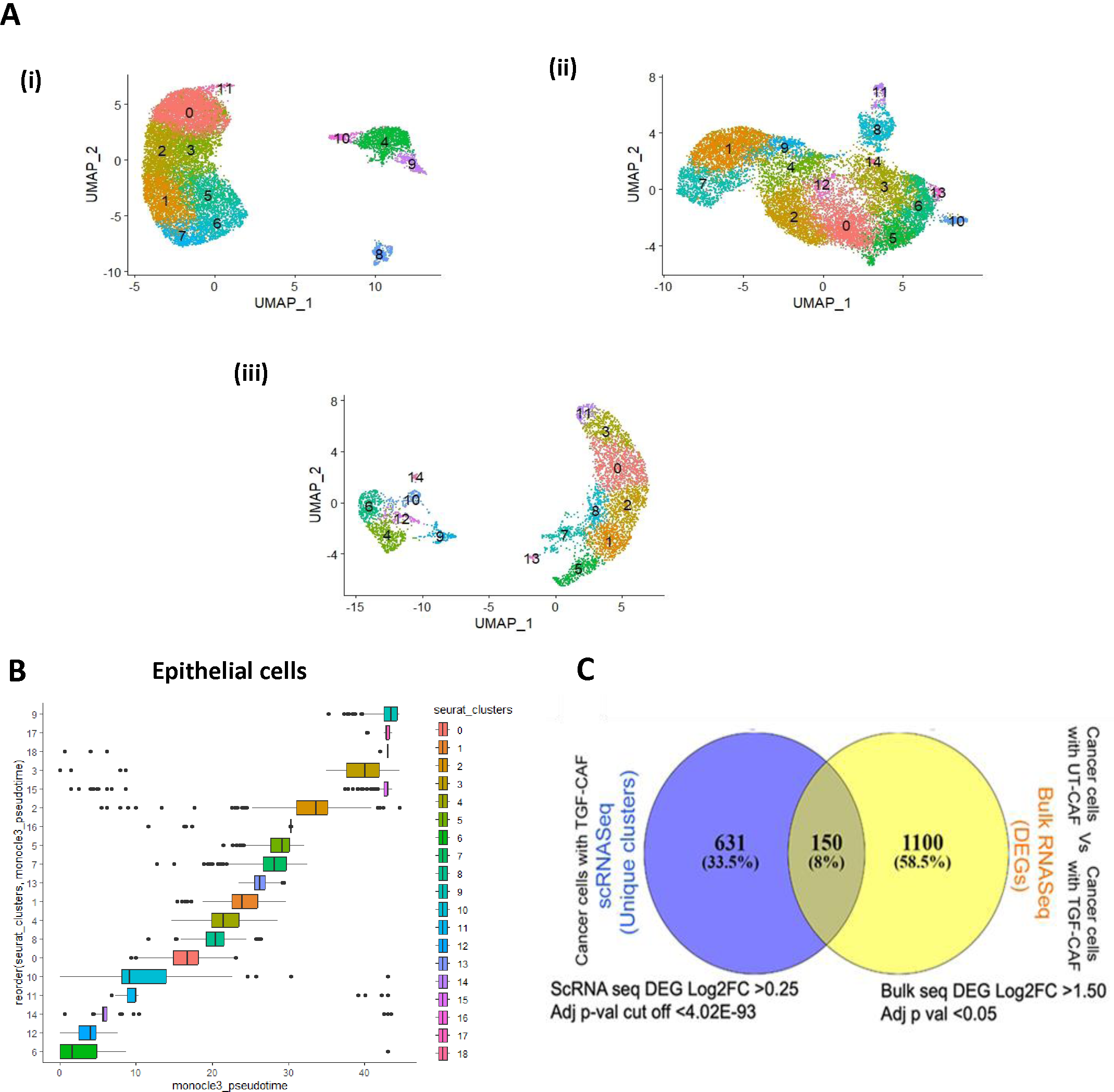
A(i-iii). Epithelial cell cluster subsets separated from the co-cultured UMAP clusters for all 3 conditions. B. Boxplot showing distribution of Monocle3 Pseudotime for each cluster of merged Cancer cell conditions. C. Venn diagram showing overlapping genes between bulk RNAseq DEGs and scRNAseq DEGs of epithelial cells co-cultured with TGF-CAF for 4 days.

**S8:**
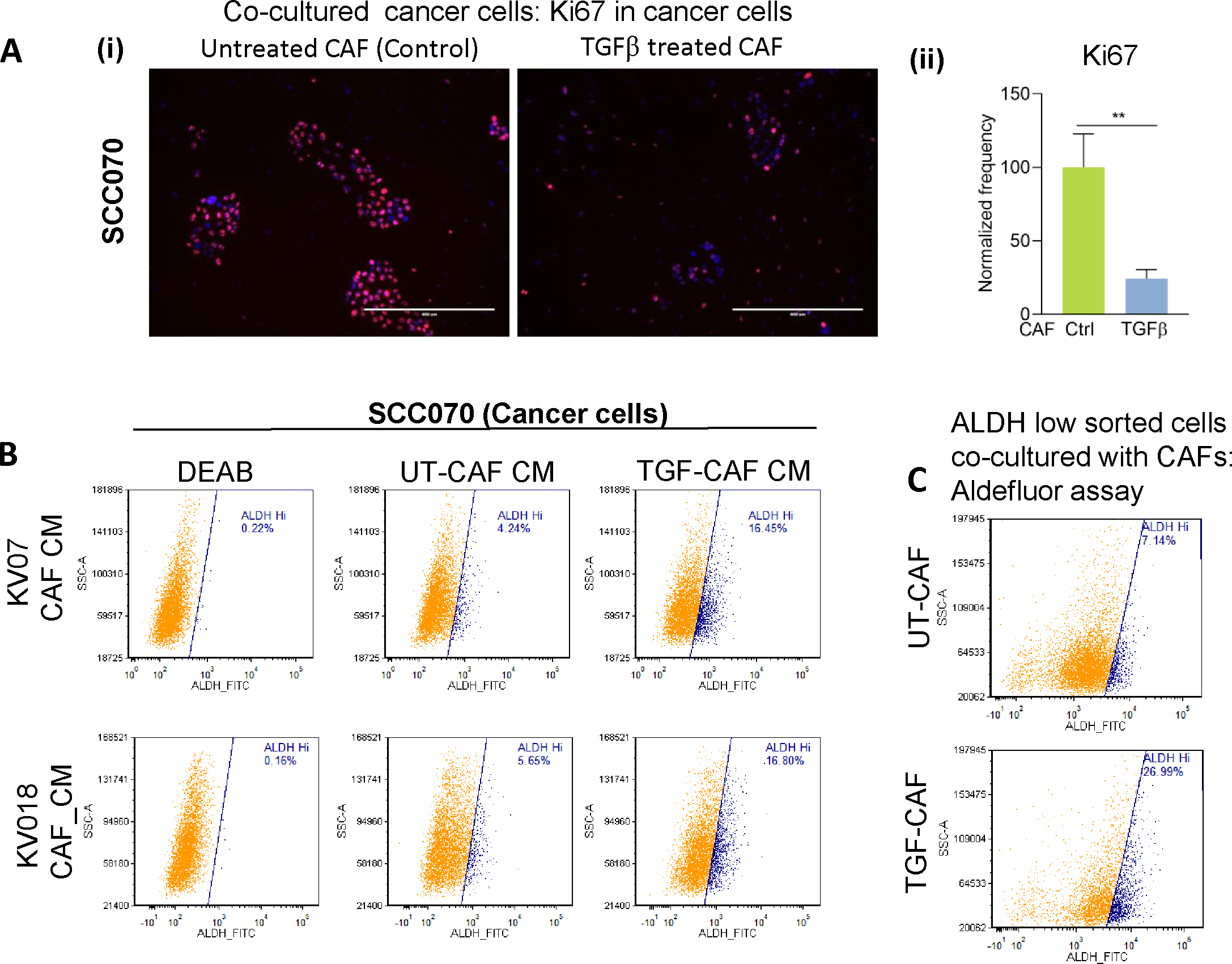
A. (i) Representative IF images of cancer cells were stained for a proliferation marker, Ki67 upon co-culture with UT-CAF and TGF-CAF for 4 days. Nuclei were stained with DAPI. Cancer cells co-cultured with UT-CAF were used as control. (ii) Frequency of Ki67 positive cells were quantified using imageJ software. Paired student t-tests were calculated. B. FACS plot showing the frequency of ALDH positive cancer cells upon exposure to UT-CAF and TGF-CAF conditioned media for 48 hrs. DEAB was used as experimental control and cancer cells exposed to UT-CAF CM was used as biological control. C. FACS plot showing the plasticity of ALDH-low cancer cells when exposed to TGF-CAF converted to ALDH-high cells. ALDH-low cancer cells co-cultured with UT-CAF were used as a control. Paired student t-test was calculated. *P<0.05, **P<0.01, ***P<0.001, ****P<0.0001 .

**S9:**
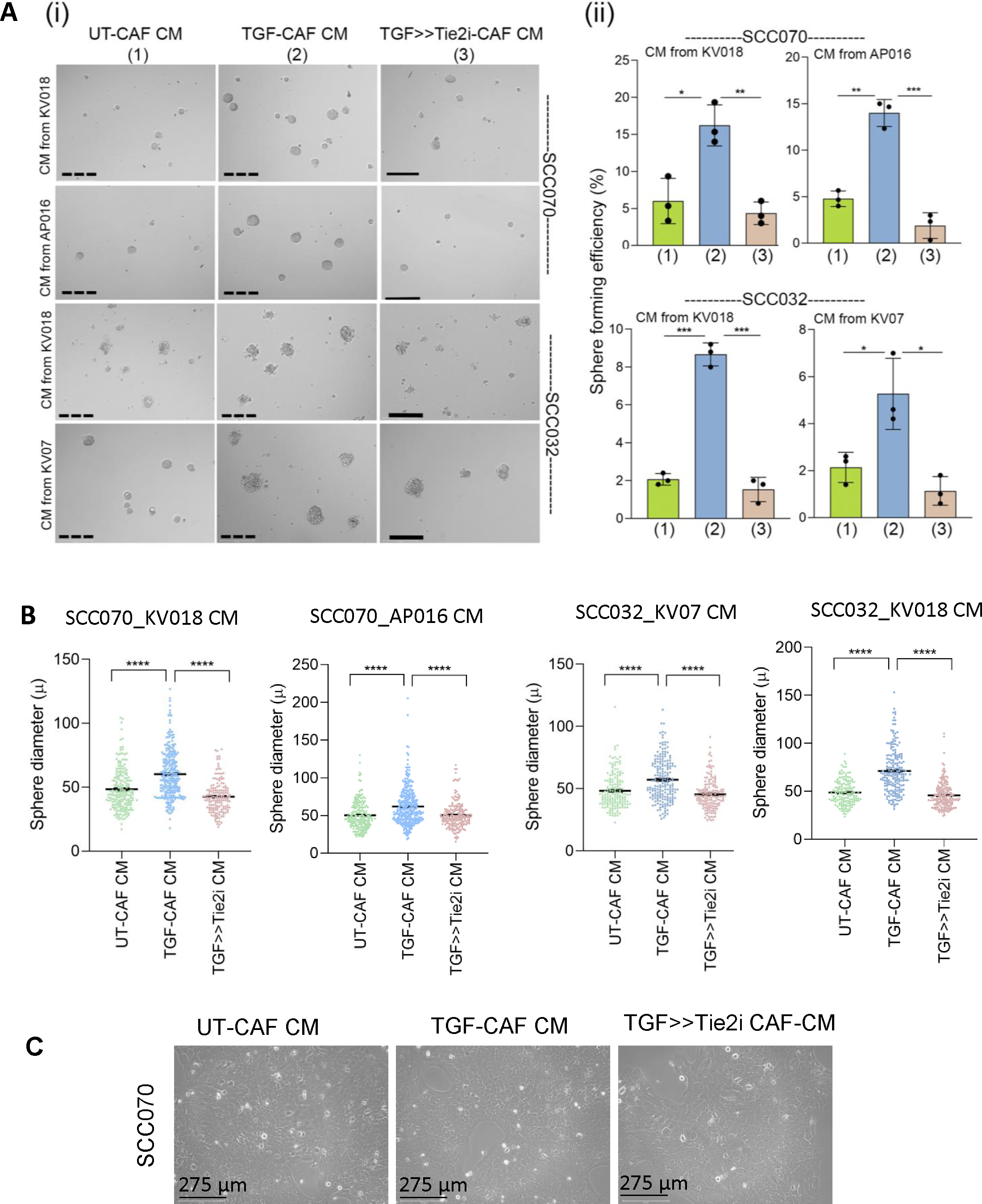
A. (i) Representative images of 3D spheroids of SCC070 cell line exposed to UT-CAF, TGF-CAF and TGF>>Tie2i-CAF conditioned media followed by testing in spheroid formation assay. Spheroids size was quantified using ImageJ. Spheroids of <60 µm diameter were excluded from study. (ii) Sphere forming frequency were calculated for the same. B. Dot plot revealing generated sphere diameter of SCC070 cells exposed to UT-CAF, TGF-CAF and TGF>>Tie2i CAF conditioned media. Cancer cells were plated in complete media for 24 hrs and media was replaced with CAF CM and kept for 48 hrs, harvested and checked for sphere forming ability up to day7. Growth factors were replenished every alternate day. Sphere size was measured and quantified using ImageJ. Statistical difference was found, as calculated by Mann-Whitney test. C. Representative phase contrast images of cancer cells exposed to UT-CAF, TGF-CAF and TGF>>Tie2i CAF conditioned media in monolayer culture. Scale bars, 275 µm *P<0.05, **P<0.01, ***P<0.001, ****P<0.0001 .

**S10:**
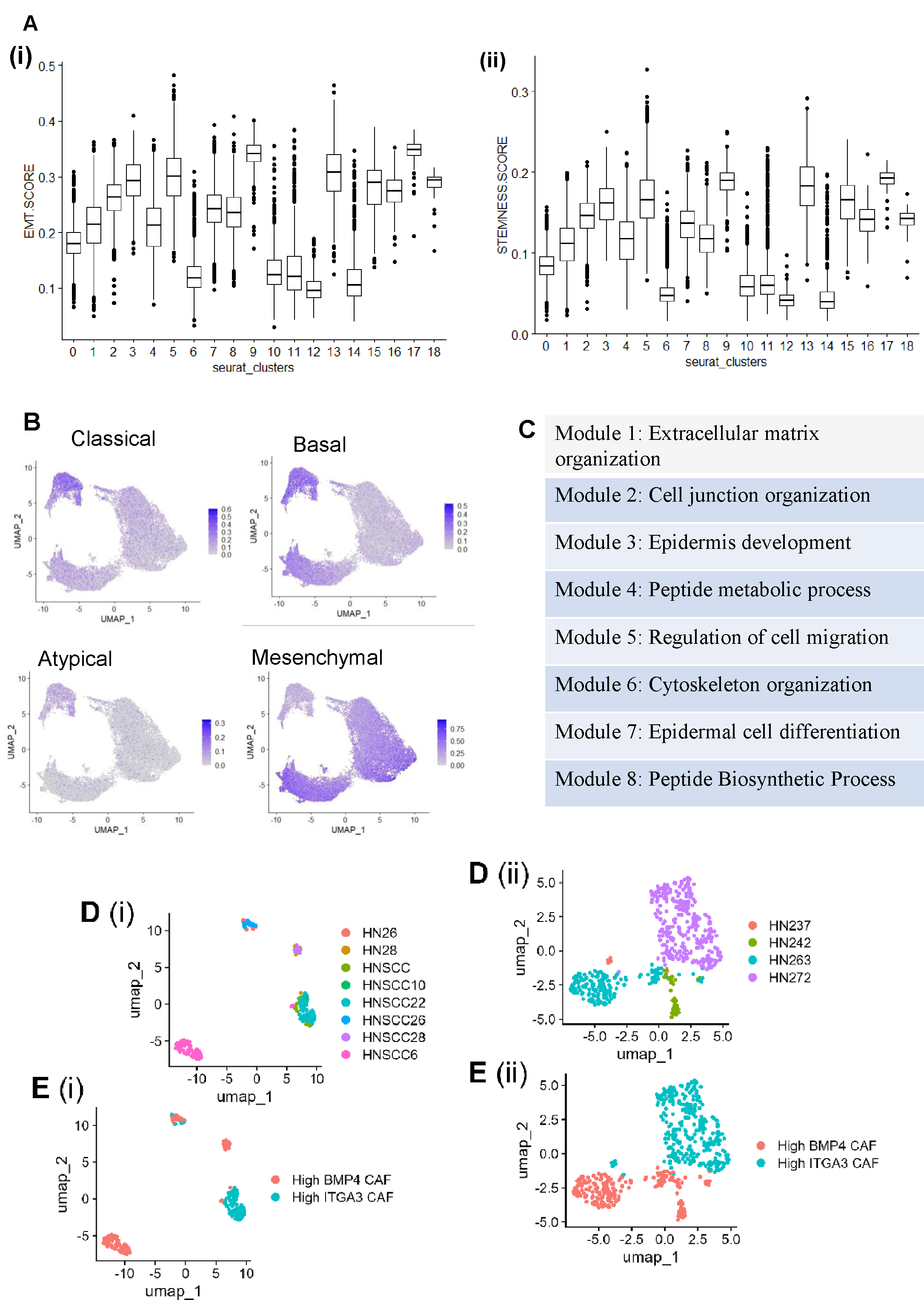
A. (i-ii) Box plot showing the distribution of stemness and EMT (Epithelial to Mesenchymal Transition) signature AUC score across different clusters of merged cancer cell subset from 3 conditions. AUC score measurement was performed by the R tool ‘AUCell’ . B. Feature plot showing the distribution of classical, basal, atypical and mesenchymal signature AUC score across different clusters of merged cancer cell subset from 3 conditions. AUC score measurement was performed by the R tool ‘AUCell’. C. Annotations of co-regulatory gene modules from GO-BP. D. (i - ii) UMAP plots of cancer cells from the classified patients, colored based on patient-wise sample-origin, or E. (i - ii) based on assigned grouping. *P<0.05, **P<0.01, ***P<0.001, ****P<0.0001

**S11:**
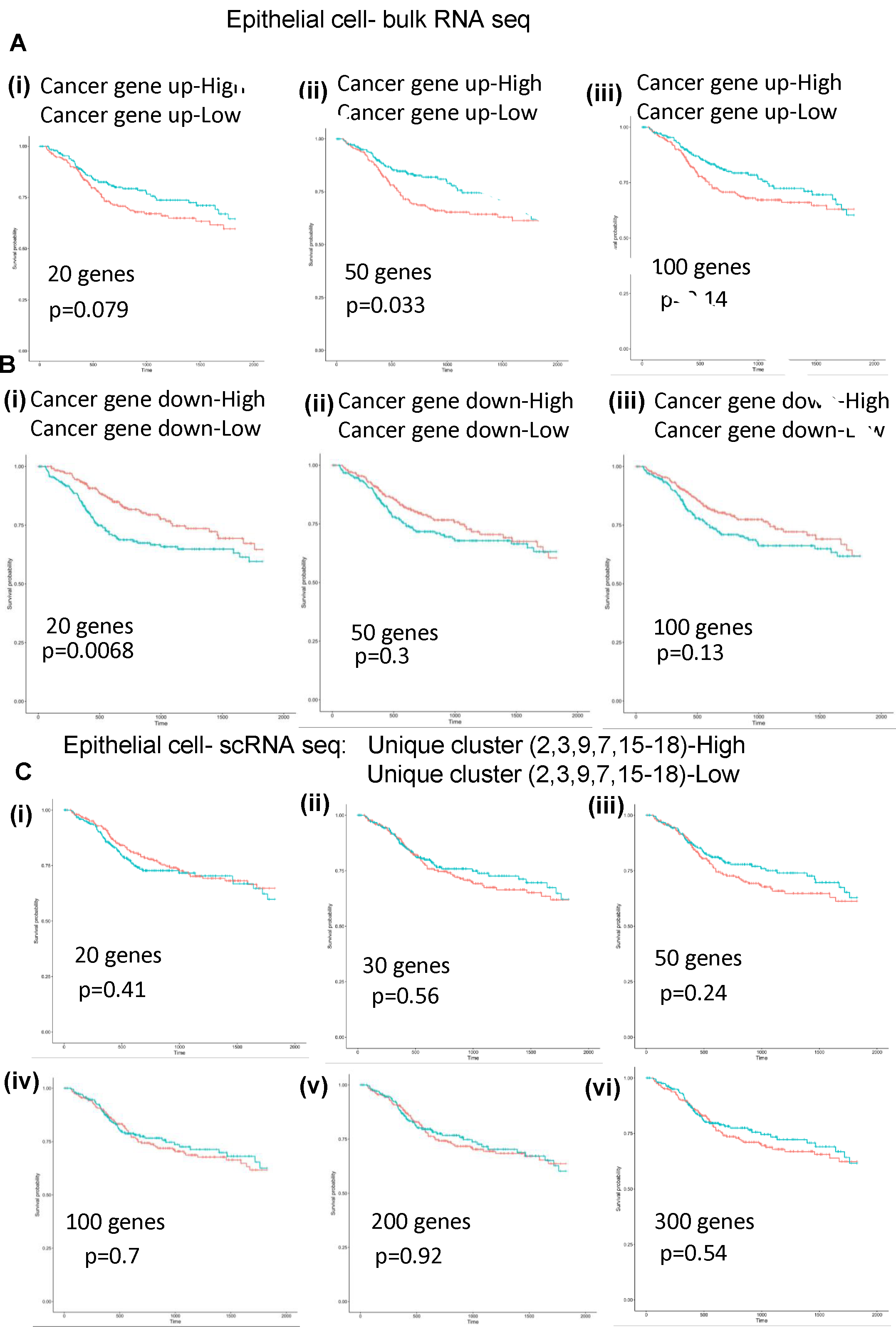
A-B. kaplan meier plot showing survival probability of HNSC patients harboring gene expression of cancer cells co-cultured with TGF-CAF. Top 20, 50 and 100 upregulated and downregulated genes were taken for single sample gene set enrichment analysis (ssGSEA). C. i-vi. Survival probability of TCGA HNSC patients expressing Top 20,30,50,100,200,300 degs of exclusive cluster (2,3,9,7,15-18) of cancer cells. Red line and blue line indicates upregulation and downregulation of specific gene signatures respectively.

